# Analgesic targets identified in mouse sensory neuron somata and terminal pain translatomes

**DOI:** 10.1101/2024.01.11.575033

**Authors:** M. Ali Bangash, Cankut Cubuk, Federico Iseppon, Rayan Haroun, Ana P. Luiz, Manuel Arcangeletti, Samuel J. Gossage, Sonia Santana-Varela, James J. Cox, Myles J. Lewis, John N. Wood, Jing Zhao

## Abstract

The relationship between transcription and protein expression is complex. We identified polysome-associated RNA transcripts in the somata and central terminals of mouse sensory neurons in control, painful (+ Nerve Growth Factor (NGF)) and pain-free conditions (Nav1.7 null mice). The majority (98%) of translated transcripts are shared between male and female mice in both the somata and terminals. Some transcripts are highly enriched in the somata or terminals. Changes in the translatome in painful and pain-free conditions include novel and known regulators of pain pathways. Antisense knockdown of selected somatic and terminal polysome-associated transcripts that correlate with pain states diminished pain behaviour. Terminal-enriched transcripts encoding synaptic proteins (e.g. Synaptotagmin), non-coding RNAs, transcription factors (e.g. Znf431), proteins associated with trans-synaptic trafficking (HoxC9), GABA generating enzymes (Gad1 and Gad2) and neuropeptides (Penk). Thus, central terminal translation may well be a significant regulatory locus for peripheral input from sensory neurons.

## Introduction

Sensory neurons are essential to drive almost all pain conditions and so their biology is of keen interest for analgesic drug development. Sensory neuron-specific transcripts,^1^ mRNA transcriptional profiles^2,3^ and proteomic studies^4,5^ have been used to examine sensory neurons in control and pain states. Distinct rather than common sets of transcriptional responses are hallmarks of different pain stimuli^2^ suggesting that a number of mechanisms drive pain from the periphery. RNA-Sequencing (RNA-seq) is often used to characterise sensory neurons but may miss low abundance RNAs.^6^ In addition, transcription is not necessarily coupled to protein translation and mRNA levels are not sufficient to predict protein concentrations.^7^ In cells as large as sensory neurons, with processes of up to a meter in length in humans, protein synthesis does not only occur in somata. Axonal protein synthesis has been observed^8^ whilst presynaptic protein synthesis in CNS neurons has been causally related to neurotransmitter release.^9^ The control of mRNA transport from soma to terminals is complex and incompletely understood.^10^

The central terminals of sensory neurons are a key regulatory site in pain pathways, as exemplified by Nav1.7 null mutant mice, where deficits in the central terminals leads to loss of neurotransmitter release resulting in a pain-free phenotype.^11,12^ Thus defining which proteins are selectively translated in sensory neuron terminals is potentially important and provides information that is not revealed by transcriptional analysis. In this study, we have used the technique of immunoprecipitating polysomes that are actively engaged in protein synthesis, so that we can identify ribosome-associated mRNAs in dorsal root ganglion (DRG) sensory neurons. Translating ribosome affinity purification (TRAP) exploits an epitope tagged (eGFP-L10) ribosomal protein as a target for antibodies that enable translating polysomes to be isolated.^13,14^ This approach has been used to examine somatic translatomes in sensory neurons in neuronal injury^15^ and neuropathic pain.^16^ These studies have also shown potential sexual dimorphism in prostaglandin signalling from somatic sensory translatomes.^17^

We modified the somatic-TRAP method (see methods) to enable specific enrichment and isolation of ribosome-bound actively translating mRNAs from both the somata and the central terminals of sensory neurons. We coupled the TRAP method with a DRG-specific Cre recombinase, using Advillin-Cre to enable DRG neuron-specific eGFP tagging of sensory neuron ribosomes. This enables isolation of actively translating mRNAs from both sensory neuronal somata and central terminals. We asked whether the translatome was gender specific, and if it changed in painful or pain free conditions. We used Nerve Growth Factor (NGF) to lower pain thresholds and the Nav1.7 knockout mouse as a pain-free mouse model to examine pain related alterations in translated proteins. We examined some identified pain-related transcripts using antisense oligonucleotides to block translation to test any significant role in pain behaviour. Here we describe the somatic and terminal translatome of mice and define known and novel analgesic targets involved in pain pathways identified with TRAP technology.

## Results

The technology employed to define ribosome-associated transcripts in the somata and terminals of sensory neurons is described in Figure 1. In order to isolate mRNAs translated in soma and central terminals of DRG neurons, we used the eGFP-L10 line,^18^ in which Cre-mediated recombination activated expression of the 60S ribosomal subunit, L10a (RPL10a) tagged to eGFP. We crossed the eGFP-L10a line with the Advillin-Cre line which expresses Cre in all DRG sensory neurons (Fig 1A), permanently labelling the ribosomes in both somata and central terminals with GFP.^19^ We sought to visualise tagged ribosomes using immunocytochemistry with GFP antibodies, detecting GFP in both the DRG soma and central terminals in the spinal cord (Fig 1B), confirming TRAP can label ribosomes in central DRG synapses. We confirmed the somatic ribosomal labelling with GFP using RT-PCR (Fig 1Ci) and immunoblotting using GFP antibodies (Fig 1Cii). The mRNA bound to ribosomes represents a very small fraction of total mRNA especially so in the central terminals and GFP-L10a levels from spinal cord were thus below the detection level of immunoblotting (Fig 1Cii), whilst the lack of a RT-PCR signal in terminal samples suggests that the ribosomal subunits are not themselves translated in the terminals but originate in the soma.

**Figure 1:**
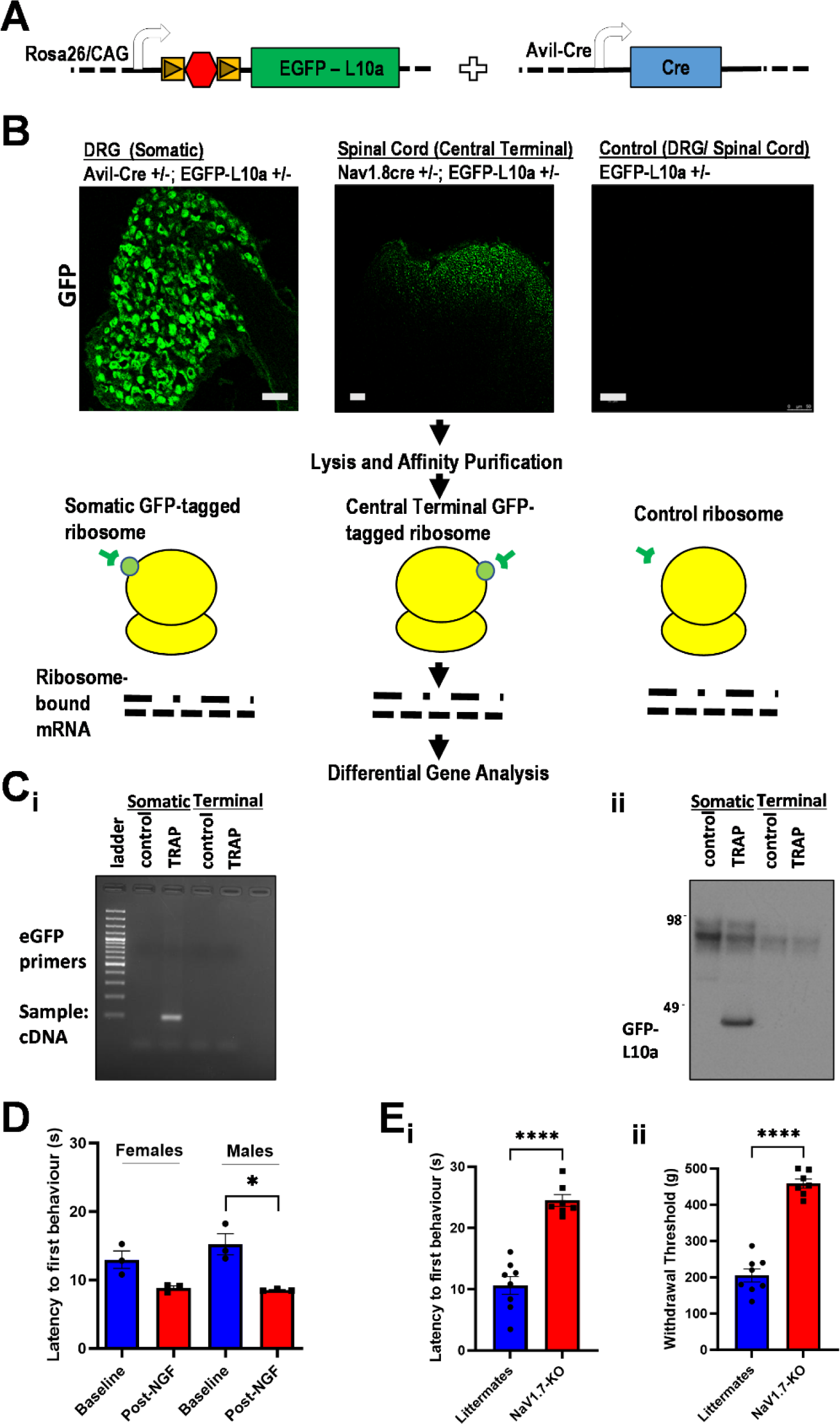
TRAP Strategy for polysomes isolation and sequencing. **A** Generation of Adv-Cre-eGFPL10 mice. eGFP L10 mice with an upstream floxed stop sequence were mated with Adv-Cre mice to generate DRG specific GFP-tagging of ribosomes. **B** Somatic and Central Terminal TRAP Strategy: Immunocytochemistry of DRG and Spinal Cord slices showing GFP-tagged ribosomal expression using polyclonal anti-GFP antibody. Scale Bar: 50μm. Tagged ribosomes were affinity purified using monoclonal anti-GFP antibodies before DGE analysis. **C** RT-PCR detection (**i**) and immunoblotting (**ii**) of GFP-tagged ribosomes from somatic and central terminal lysates. **D** NGF-evoked increased pain sensitivity was measured using the Hargreaves’ test in both male and female mice. **E** The acute pain-free status of Advillin-Cre Nav1.7 null mice was confirmed with measurements of thermal (**i**) and mechanical (**ii**) thresholds. Mean latencies and withdrawal thresholds in **D** and **E** were compared using unpaired t-test. * = p<0.05; **** = p<0.0001.

In order to assess the level of background mRNA non-specifically binding to immunoglobulins, beads or protein L, we examined the signal from eGFP-L10a lines without Cre, detecting no GFP signal (Fig1B, Ci, and Cii) and detecting negligible amounts of total mRNA immunoprecipitated from these Cre-negative samples. Therefore, the immunoprecipitation of GFP-tagged ribosomal-mRNA complexes from dissected DRG and spinal cord seems to reflect polysome-associated mRNA in these samples.

We needed to generate mice in different pain states, and we selected Advillin-Cre floxed Nav1.7 null mutant mice as pain free examples,^12^ as well as normal mice treated with NGF as a model for inflammatory pain (Figure 1D, E)^20^.

### Sex-specific translatomes?

We analysed somatic ribosome bound mRNAs (the translatome) from dissected DRGs from Adv-Cre-GFPL10a mice using RNA-seq. We compared male Adv-Cre; eGFPL10a samples directly with female samples (Fig 2A). The alternative method of normalizing the TRAP IP (translatome) to the unbound supernatant (“transcriptome”)^14^ is likely to dilute the sensory neuron translatome because the unbound fraction from a heterogenous tissue such as the spinal cord or DRG will contain many non-neuronal mRNAs.

**Figure 2:**
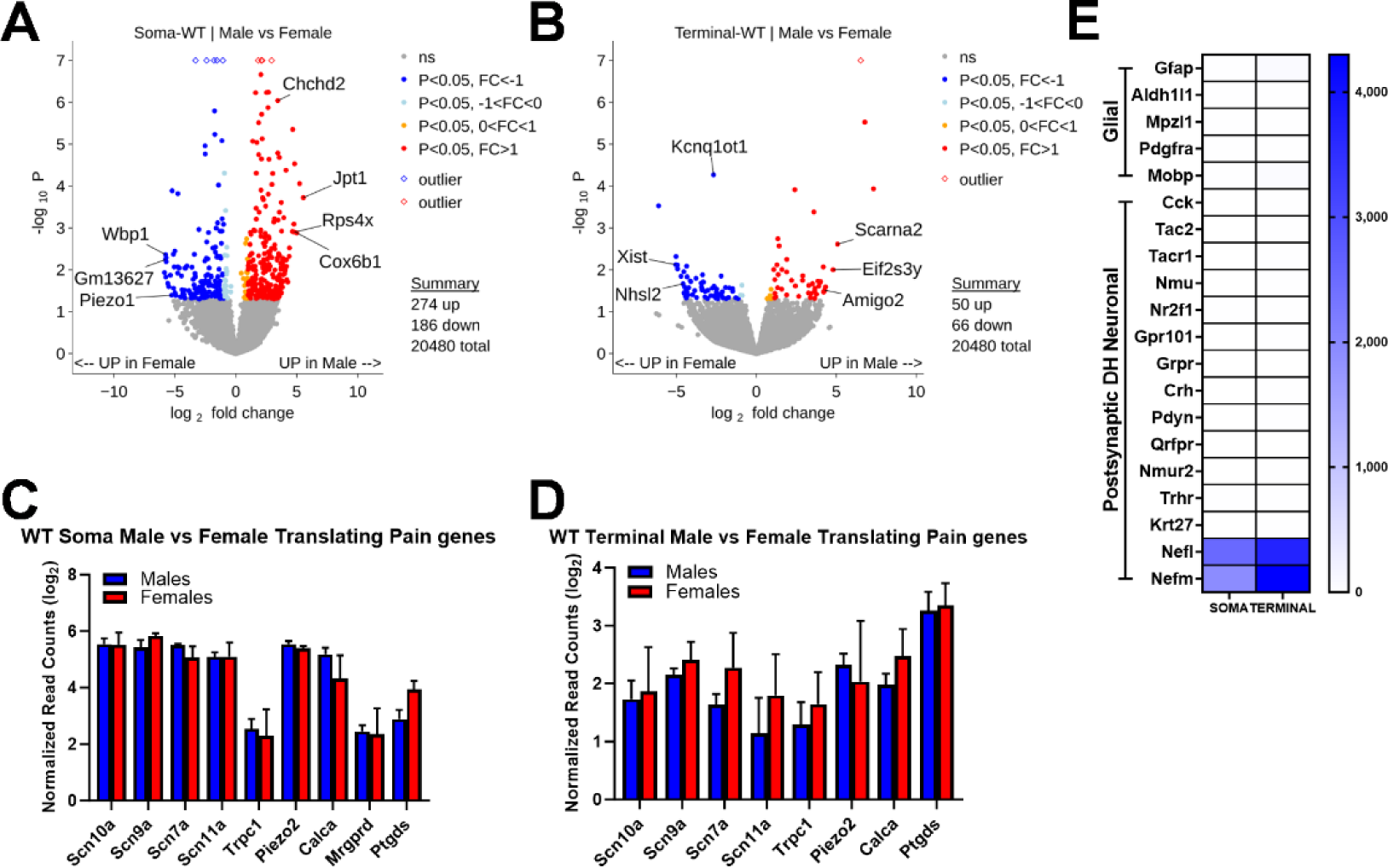
Sex differences in male and female translatomes. **A** Volcano plot showing differentially expressed translated genes in the soma between male (n=6) and female (n=6) samples. Statistical analysis by Wald test using DESeq2. Genes on the left side of the plot are upregulated in females and upregulated male genes are shown on the right side. Non-significant genes are shown in grey, significant genes (p<0.05) with log_2_ fold change < 1 with blue and significant genes with log_2_ fold change > 1 are represented with red colour. X-axis and y-axis show, log_2_ fold change and -log_10_ p-value, respectively. Complete data comprising read numbers and fold increase (log_2_) with p-values are presented in excel format in table S1. **B** Same as in **A**, but the volcano plot shows the genes expressed in the terminals. Complete data comprising read numbers and fold increase (log_2_) with p-values are presented in excel format in table S2. A volcano plot showing pooled data from somata and terminals is detailed in figure S1. **C** Bar plot comparing the expression levels of known pain genes in male (in blue) and female (in red) somata. **D** Bar plot comparing the expression levels of known pain genes in male (in blue) and female (in red) terminals. **E** Heatmap showing the Log read counts of different non neuronal genes. The expression of these genes is very low when compared to control genes (Nefl and Nefm) expressed at similar levels in soma and terminals translatomes. Normalized read counts in **C** and **D** were compared using unpaired t-test.

The potential problem of low-level contamination of RNA transcripts in our TRAP samples is to some extent obviated by the fact that concentrating on high read number altered transcripts in different protocols is likely to reflect specific immunoprecipitation profiles. We sequenced 20,480 genes from somatic male and female Adv-Cre; eGFPL10a DRGs, out of which 460 were differentially expressed (Fig 2A, B, and Tables S1, S2).

Since TRAP isolates ribosome-bound mRNAs in a cell-type specific manner, we tested the specificity of our TRAP IP by examining the expression of non-neuronal DRG genes (e.g. *GFAP*) and found them to be completely absent from IP samples (Fig 2E). This confirms the specificity of the TRAP pull down. We have listed all sequenced genes with read counts, fold changes, log2 fold changes and p-values in a series of supplementary tables that link to each of the Volcano Figures (Figure S1 and Tables S1-S9).

### Advillin-Cre; eGFPL10a somatic and terminal translatomes are shared by male and female mice

Despite the low abundance of ribosome-bound mRNA in central terminals of DRGs, we were able to successfully isolate and sequence the terminal translatome from pooled dissected spinal cords from Advillin-Cre; eGFPL10a mice (Fig. 2B). Although we sequenced a higher number of low-read number genes in the terminal compared to the somatic TRAP, this likely represented a higher background from the greater mass of spinal cord tissue.

Comparing male and female DRG sensory neuron total translatomes directly, we identified a small fraction of sex specific transcripts (116 of 20480 genes), indicating predominantly shared pain pathways including key pain genes (Fig 2C, D). We tested the specificity of our terminal IP by examining the expression of non-neuronal DRG genes (for example GFAP) and postsynaptic dorsal horn neuronal genes (for example PDYN absent in normal DRG neurons^21^) which were found to be entirely absent in our transcriptome data sets (Fig 2E).

The vast majority (>98%) of the translatome was shared between male and female mice suggesting predominantly shared pain mechanisms, including key pain genes (e.g. *Scn9a*, *Scn10a*) (Fig 2C, Tables S1, S2). These numbers are consistent with recent TRAP-seq studies^22,23^, including a study of sex differences from TRAP samples in Nav1.8+ neurons^24^. In total DRG neuronal samples we found a small increase in *Ptgds* mRNA in female mouse somata with low-read numbers (See Table S1) consistent with the data in Tavares-Ferreira *et al.*^24^

When comparing somatic male and female translatome data we found that *Piezo1*, a mechanosensitive channel, was upregulated in females although transcript levels were low. In female terminals, *Kcnq1ot1* is 8-fold enriched – this is a long ncRNA that is known to interact with chromatin.^29^ Intriguingly *Xist*, which is totally absent from somatic translatomes, is found in central terminals and enriched in female mice.^25^ This gene, crucial for X-inactivation, may associate with polysomes like other ncRNAs but its mechanisms of action remain uncertain. *Nhsl2* is a cytoplasmic calcium-binding protein 20-fold upregulated in females that is linked to Nance-Horan syndrome.^26^

In male somata, the gene *Chchd2* implicated in stress responses to low oxygen was upregulated 20-fold with high read numbers^27^ whilst *Jpot1*, which encodes a nuclear envelope protein, was 50-fold upregulated. Very interestingly, the mRNA encoding the isoform of ribosomal protein RPS4X is 30-fold enriched in the male translatome, suggesting that there is a potential for gender specific differences in ribosomal composition at the terminals.^28^

The top genes differentially upregulated in male terminals include the translational control gene *Eif2s3y*^28^ which is 30-fold upregulated in males, and is known to enhance synaptic efficacy in male but not female mice. Its overexpression is linked to autism.^29^ *Scarna2* is an ncRNA that has multiple functions in DNA repair and is 30-fold upregulated in males, whilst *Amigo2* has been implicated in synaptic function and cell-cell interactions, and is 20-fold upregulated in males.^30^

Taken together the results support the claim that terminal translatome transcripts accurately reflect transcripts present in sensory neurons but generally absent from other cell types (Table S2). The fact that major pain related genes are shared by male and female mice led us to focus on male mice to minimise animal usage.

### Distinct translatomes in somata and terminals

Although 90% of transcripts were common to terminal and soma translatomes, and those present at very low read levels potentially included some contaminating material, about 5% of transcripts were enriched either in the soma or in the terminals, (1704 genes from a total of 20,480 examined) some more than 30-fold (Figure 3 and S2, and Table S3). There were some interesting findings. For example, *Penk* mRNA encoding the enkephalins, was substantially enriched in the terminals and absent from the soma, a finding consistent with recent insights into opioid signalling within the spinal cord and regulation of pain pathways.^12^ In contrast, Calcitonin Gene Related Peptide (CGRP) genes were principally translated in the soma, perhaps because these peptides play an important role at the peripheral terminal in regulating blood flow.^31^ Voltage-gated sodium channels are a topic of interest for pain studies, and Nav1.1 was enriched in the terminals of sensory neurons, whilst Nav1.7, Nav1.8 and the key regulator of Nav1.8 expression p11 (S100A10) were substantially enriched in somatic polysomes.^32^ Enzymes associated with the production of GABA (Gad1 and Gad2) were enriched in the translatome with high read numbers.^33^ The complete data are to be found in supplementary tables and potential functions are debated in the discussion section.

**Figure 3:**
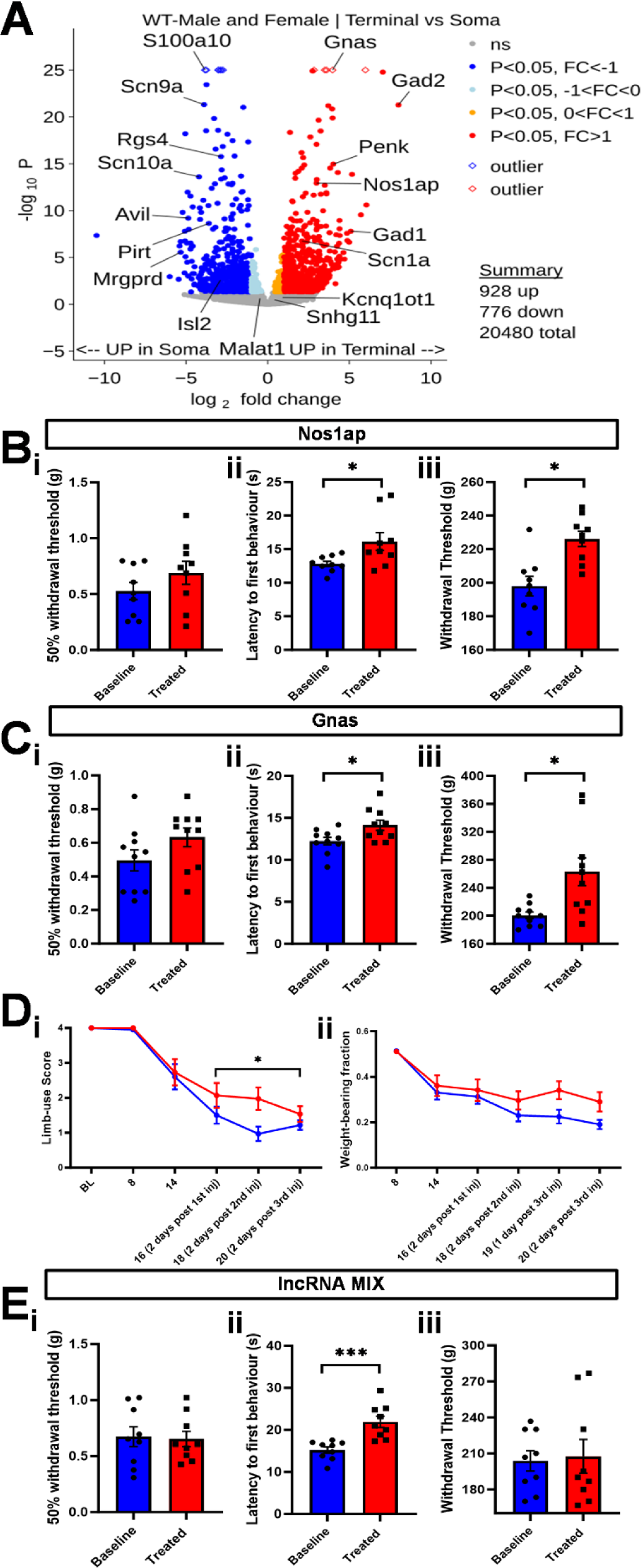
Somatic versus terminal translatomes. **A** Volcano plot showing differentially expressed translated genes between somata (n=6) and terminal (n=6) samples. Genes on the left side of the plot are upregulated in somata and upregulated terminal genes are shown on the right side. Non-significant transcripts are grey, significant genes (p<0.05) with log2 fold change < 1 with blue and significant genes with log_2_ fold change > 1 are represented with a red colour. x-axis and y-axis show, log_2_ fold change and -log_10_ p-value, respectively. Complete data comprising read numbers and fold increase (log_2_) with p-values are presented in table S3. Futher comparisons between somata and terminal genes, as well as gene enrichment pathways in both conditions, are detailed in figure S2. **B** Antisense oligonucleotide (ASO) inhibition of terminal translatome transcripts Nos1ap lowers heat and mechanical pain thresholds. Mice (n=10) were tested with the von Frey (**i**), Hargreaves’ (**ii**) and Randall Selitto (**iii**) tests before (baseline) and after ASO intrathecal injection (treated). **C** Antisense oligonucleotide (ASO) inhibition of terminal translatome transcripts Nos1ap lowers heat and mechanical pain thresholds. Mice (n=10) were tested with the von Frey (**i**), Hargreaves’ (**ii**) and Randall Selitto (**iii**) tests before (baseline) and after ASO intrathecal injection (treated). Substantially diminished pain behaviour was apparent in both **B** and **C**. Experiments using control ASOs are shown in figure S3. **D** Cancer induced bone pain is attenuated by ASO knockdown of Gnas. Limb use and weight bearing were improved with ASO knockdown. **E** ASO knockdown targeting combined lncRNAs Malat1, Kcnq1ot1, and Snhg11 leads to lowered heat pain behaviour as measured with a Hargreaves’ apparatus. Experiments silencing individual lncRNAs are shown in figure S4. Motor impairment tests for **B**, **C**, and **E** are shown in figure S5. Mean latencies and withdrawal thresholds in **B**, **C**, and **E** were compared using paired t-test. Limb scores and weight-bearing fractions in **D** were compared using restricted maximum likelihood analysis (REML). * = p<0.05; *** = p<0.001.

We investigated the role of some peripherally enriched transcripts using antisense knock-down. *Nos1ap* has been linked to neuropathic pain in global knock-out studies.^34^ It is also implicated in neuropathic pain in mice and some progress has been made in developing blockers of interactions with NOS. Our data suggest that terminal *Nos1ap* plays a role in pain induction. *Gnas* is an interesting and complex imprinted gene that encodes the alpha subunit of the adenyl cyclase activating complex Gs. Gain-of-function mutations in *GNAS* are associated with painful conditions in humans.^35^ We found a clear inhibition of thermal and mechanical acute pain with antisense oligonucleotides (ASO) directed against *Gnas* delivered intrathecally, whilst control scrambled ASOs were inactive (Figures 3B, C, and Figure S3A, B). Given the role of adenyl cyclases in inflammatory pain, this is an interesting potential target for localised pain therapies. To further check this hypothesis, we tested the translation knockdown of *Gnas* in a mouse model of cancer-induced bone pain using three intrathecal injections of ASOs against *Gnas* (Figure 3D). The knockdown of this gene resulted in a modest reduction of pain-like behaviour associated with this model, as assessed by limb-use scoring and weight-bearing. Statistical analyses indicated that the knockdown of *Gnas* expression slowed down the reduction of the limb-use significantly compared to control mice (Figure 3Di) (REstricted Maximum Likelihood (REML) analysis, p-value=0.0486). While the improvement of weight-bearing following *Gnas* knockdown was apparent, it failed to reach an 0.05 level of statistical significance (REML, p-value= 0.0537, Figure 3Dii). It is likely that our antisense protocols will diminish both transcript levels as well as the translation of candidate target genes. Nevertheless, if there is less mRNA to translate, this approach still relates to the role of the translatome in regulating pain pathways.

The TRAP RNA-seq data not only identified protein-coding mRNAs but also a significant number of non-coding RNAs that were associated with the sensory neuron ribosomes^36^. This is consistent with previous polysome profiling data where it was found that the majority (70%) of cytoplasmic-expressed long non-coding (lnc) RNAs have more than half of their cytoplasmic copies associated with polysomal fractions^37^. We focused on three high read number ncRNAs associated with sensory neuron polysomes. *Malat1*^38^, *Kcnq1ot1*^39^ and *Snhg11*^40^ and found that antisense knockdown could lower heat pain thresholds when all three ncRNAS were targeted, without any effect shown from the scrambled control ASOs (Figure 3E, Figure S3C). However, individually targeted ASOs did not have any statistically significant effect (Figure S4), suggesting a minor contribution to heat thresholds of the tested ncRNAs. Whether the sensory neuron polysome-associated lncRNAs identified here actively participate in regulating translation, or have other ribosomal functions, in different pain states merits further investigation.

### NGF-induced pain states alter translatome activity

In order to address potential alterations in the translatome int pain states, we used the well characterised inflammatory mediator NGF to sensitise mouse tissues globally (Figure 1D). We then used TRAP technology to compare the polysome-associated transcripts in an NGF-evoked pain state with normal mice (Tables S4, S5). A number of studies have examined translational alterations in NGF-evoked pain states in mice, but these events do not seem to be mirrored in the polysome-associated transcripts that we identified^41^.

Nerve growth factor acting through TrkA is known to sensitise pain pathways at the level of sensory neuron activation. We examined the alterations in the soma and terminals comparing NGF-treated samples with control samples. We found 268 genes of 20,480 examined were dysregulated in the soma, whilst more changes (369 transcripts) were altered in the terminal from the 20,480 examined (Figure 4).

**Figure 4:**
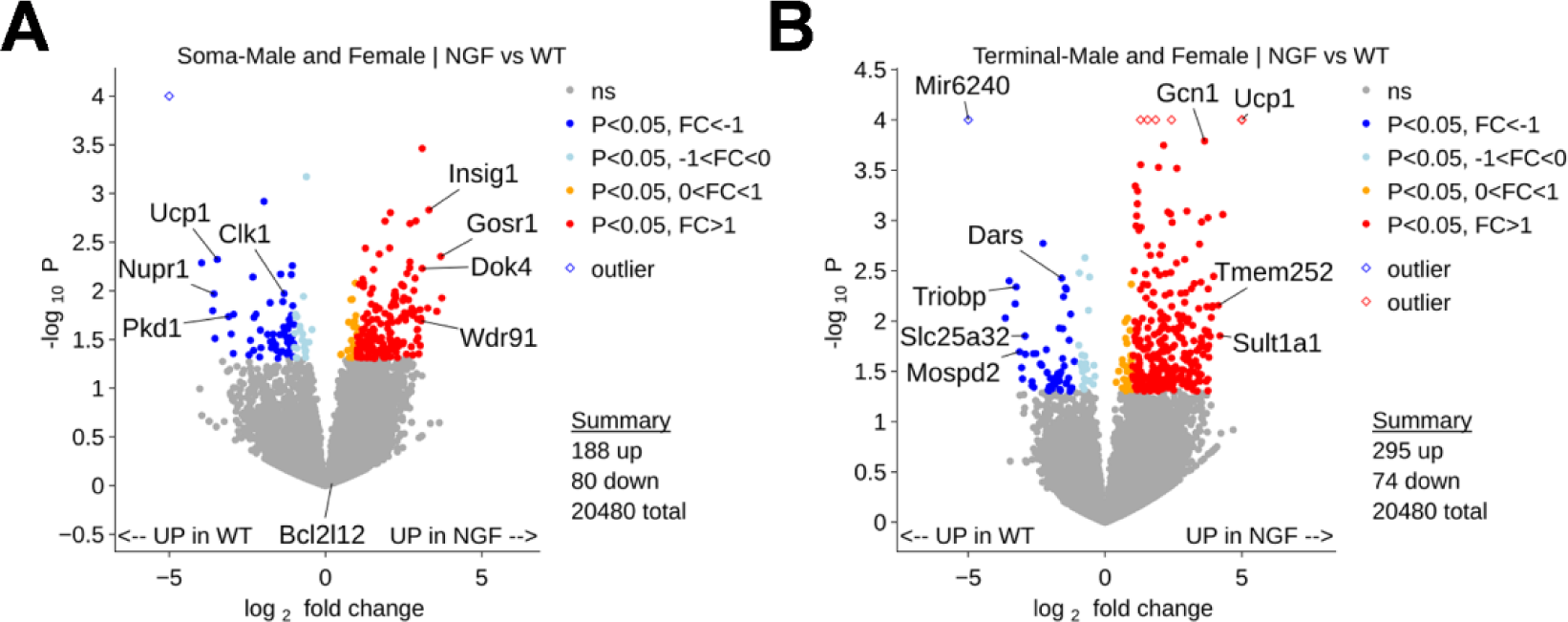
NGF-induced transcripts in somatic and terminal translatomes. Volcano plots showing differentially expressed translated genes between NGF enriched (n=6) and control (n=6) samples in the somata (**A**) and terminals (**B**). DESeq2 was used for statistical analysis. Genes on the left side of the plot are down regulated by NGF and NGF upregulated genes are shown on the right side. Non-significant genes are in grey, significant genes (p<0.05) with log_2_ fold change < 1 with blue and significant genes with log_2_ fold change > 1 are represented with red colour. x-axis and y-axis show, log_2_ fold change and -log_10_ p-value, respectively. Complete data comprising read numbers and fold increase (with log_2_) with p-values are presented in excel format in tables S4 and S5. Histograms detailing the gene enrichment pathways in both conditions are shown in figure S6.

A range of mRNAs were upregulated in the somatic translatome on NGF treatment (Figure 4A). These included mitochondrial proteins that could be involved in increased metabolic activity in activated sensory neurons, *Unc79* which encodes the mouse homologue of UNC79,^42^ which is a subunit of the sodium ion leak channel NALCN, and *Dok4*, a transmembrane tyrosine kinase receptor involved in regulating neurite outgrowth during nervous system development^43^.

In terminals (Figure 4B) we found that *Gcn1*, whose encoded protein enhances translation by removing stalled ribosomes and is associated with polysomes, was strongly upregulated consistent with increased protein synthesis.^44^ Other enhanced transcripts included phenol sulfotransferase *Sult1a1,*^45^ whose human homologue is highly inducible by dopamine and is part of a family of proteins thought to protect neurons from neurotoxicity,^46^ as well as*Tmem252* thought to play a possible role in kidney function.^47^ Bioinformatic analysis links enhanced glutamatergic synapse activity as well as PI3 Kinase Akt signalling activity with the terminals to NGF-induced enhanced pain states – in agreement with such enhanced activity found in the spinal cord in chronic pain (Figure S6)^48^.

### Pain-free Nav1.7 null mouse translatomes

We examined the alteration in translatomes in the pain free Nav1.7 null mouse (Figure 5, Tables S6, S7). Amongst genes downregulated in the somatic translatome are several that were not detected in microarray analysis of Nav1.7 null DRG^49^. Genes shown in Figure 5 could be potential mediators in pain pathways. 132 genes were down regulated in soma and 557 were lower in terminals of pan-DRG pain-free Nav1.7 null mice.

**Figure 5:**
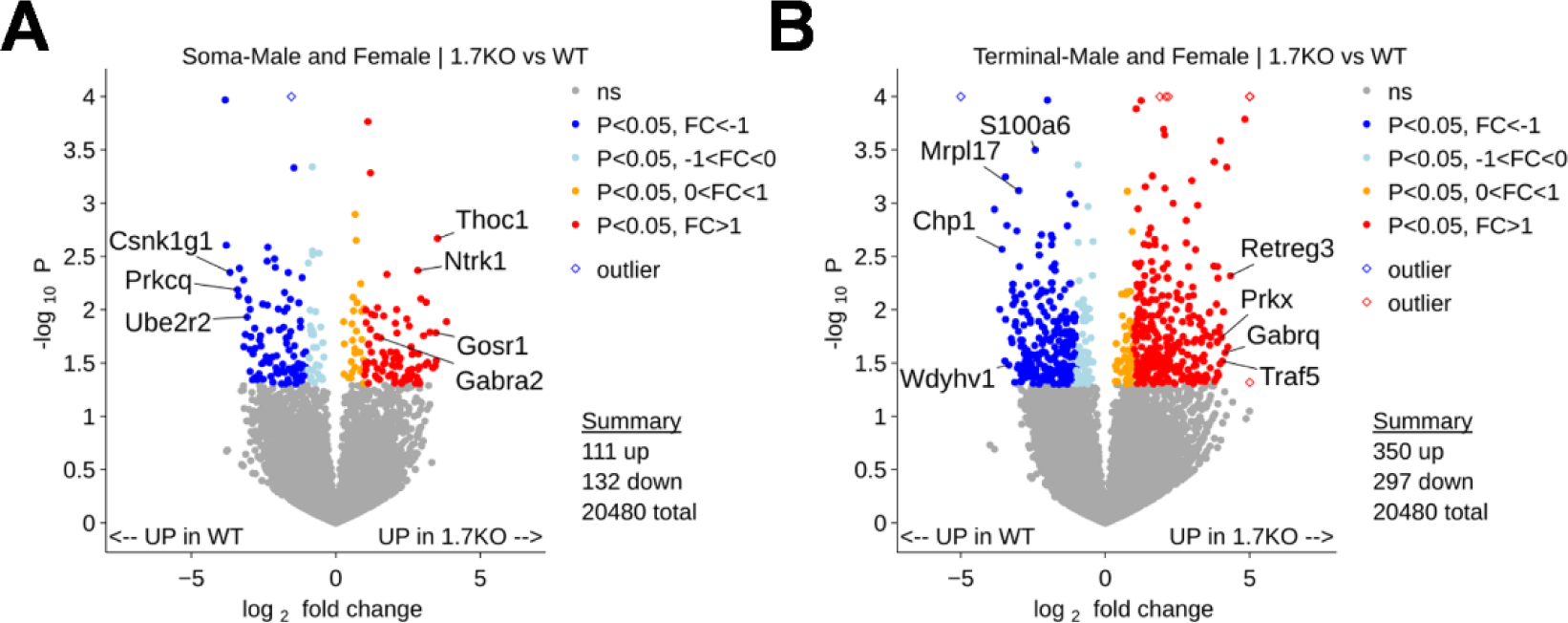
Altered translatomes in Nav1.7 null, pain-free mice soma and terminals. Volcano plots showing differentially expressed translated genes between Nav1.7 null (n=6) and control (n=6) samples in the somata (**A**) and terminals (**B**). Genes on the left side of the plot are down regulated in Nav1.7 nulls and upregulated genes are shown on the right side. Non-significant genes are in grey, significant genes (p<0.05) with log_2_ fold change < 1 with blue and significant genes with log_2_ fold change > 1 are represented with red colour. X-axis and y-axis show, log_2_ fold change and -log_10_ p-value, respectively. Complete data comprising read numbers and fold increase (log_2_) with p-values are presented in excel format in tables S6 and S7. Histograms detailing the gene enrichment pathways in both conditions are shown in figure S7.

Amongst the genes downregulated in Nav1.7 null somata (Figure 5A), the *Clock* gene is a bHLH transcription factor that controls the expression of the Per genes that in DRG regulates neuronal excitability with a circadian rhythm.^50^ *Mrgprd* is a GPCR that is activated by enkephalins and other ligands.^51^ Protein kinase C theta (*Prkcq*) is a calcium-independent serine-threonine kinase involved in neurotransmitter release.^52^ Tyrosine hydroxylase (*Th*) generates dopamine from tyrosine that then gives rise to catecholamines all of which are implicated in pain regulation. Casein kinase 1 gamma 1 (*Csnk1g1*) phosphorylates acidic proteins on serine and threonine and has been linked to epilepsy^53^.

Within the terminals (Figure 5B), transcripts reduced in the Nav1.7 null mice translatome include *Ppp6r1*, a phosphatase that may be involved in NF-kappa B signalling.^54^ *Mepce* is involved in RNA methylation and enhances Pol2 dependent transcription.^55^

*Tssc4* is a tumour suppressor implicated in a large range of pathologies.^56^ *Chp1* is a key component of the Na/H exchanger and is a calcium binding EF hand protein.^57^ Inflammatory pain as well as intracellular pH control are linked to these proteins. *Hhipl1* is a hedgehog interacting protein linked to morphogenesis and differentiation.^58^

Bioinformatic analysis shows diminished glutamatergic and dopaminergic synapse activity in Nav1.7 nulls, and oxidative phosphorylation genes are also downregulated in both somata and terminals in the pain free state. This fits with less sensory neuron activity in the absence of nociceptive input.

### Identifying translated proteins that correlate with pain states

Now that we had the repertoires of translated genes in pain-free and painful states from somata and terminals of DRG sensory neurons, we were able to focus on translated genes that correlate strongly with enhanced pain and examine their potential significance using antisense knock-down behavioural assays (Figure 6).

**Figure 6:**
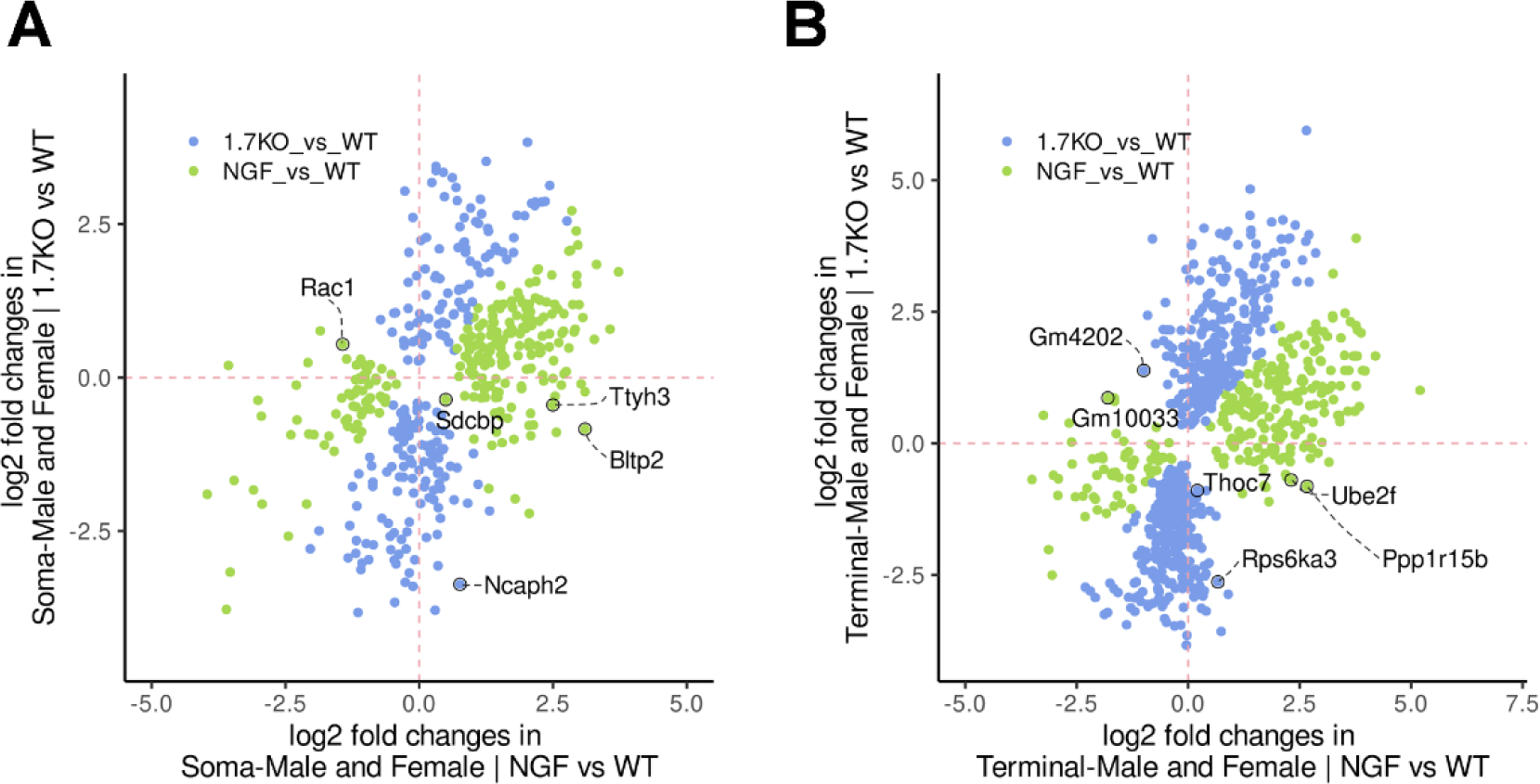
Differences in gene expression between NGF-treated and NaV1.7 null mice soma and terminals. Scatter plots showing differentially expressed translated genes between Nav1.7 null (n=6) and NGF treated (n=6) samples in the somata (**A**) and terminals (**B**). Non-significant genes are not showed; significant genes (p<0.05) in the NGF vs WT comparison are shown in green, whereas significant genes (p<0.05) in the 1.7 null vs WT comparison are shown in blue. X-axis and y-axis show log_2_ fold change for the two comparisons. Complete data comprising read numbers and fold increase (log_2_) with p-values are presented in excel format in tables S8 and S9. Histograms detailing the gene enrichment pathways in both conditions are shown in figure S8.

We examined the relative expression of promising candidate genes in pain states, in the pain free mouse and the wild type controls to select potential targets for antisense experiments. Volcano plots are shown in Figure 4, 5, and 6, where a number of potential targets are identified.

We compared expression of some of the most promising candidate genes in the somata of mice as shown in Figure 7A, where the relative expression is colour coded. We found some transcripts were present at very low levels in both pain free and normal mice but highly expressed with NGF (e.g. *Wdr91*^59^, *Cav2*), whilst others showed a gradation of increased expression that correlated with enhanced pain states (e.g. *Sdcbp*^60^, *Rgs4*^61^) (Figure 7B). *Sdcbp* or syntenin is associated with kainate receptor expression whilst the *Necab* family of genes are neuronal specific^60^ and *Necab2* is associated with mGlu receptors linked to autism and neurodegeneration^62,63^. Sensory neuron glutamate receptors have been implicated in altered pain states. *Ttyh3*, a chloride channel, has been implicated in epilepsy, chronic pain, and viral infections.^64^ Related channels have been reported to form Ca^2+^-and cell volume-regulated anion channels structurally distinct from any characterized protein family with potential roles in cell adhesion, migration, and developmental signalling. *Wdr91* is implicated in neurodegeneration and lysosome function.^59^ *Psme3* activates the proteosome,^65^ whilst *Cav2* is a caveolin-like molecule of unknown function. *Mtmr3* or Myotubularin-related protein regulates the cell cycle.^66^ *Ncaph2* is implicated in cognitive decline and AD,^67^ whilst *Rgs4* is linked to schizophrenia and G protein function. *Rac1* is a small GTPase linked to the inflammasome^68^ and *Uhrf1bp11* is a lipid transfer bridge.^69^

**Figure 7:**
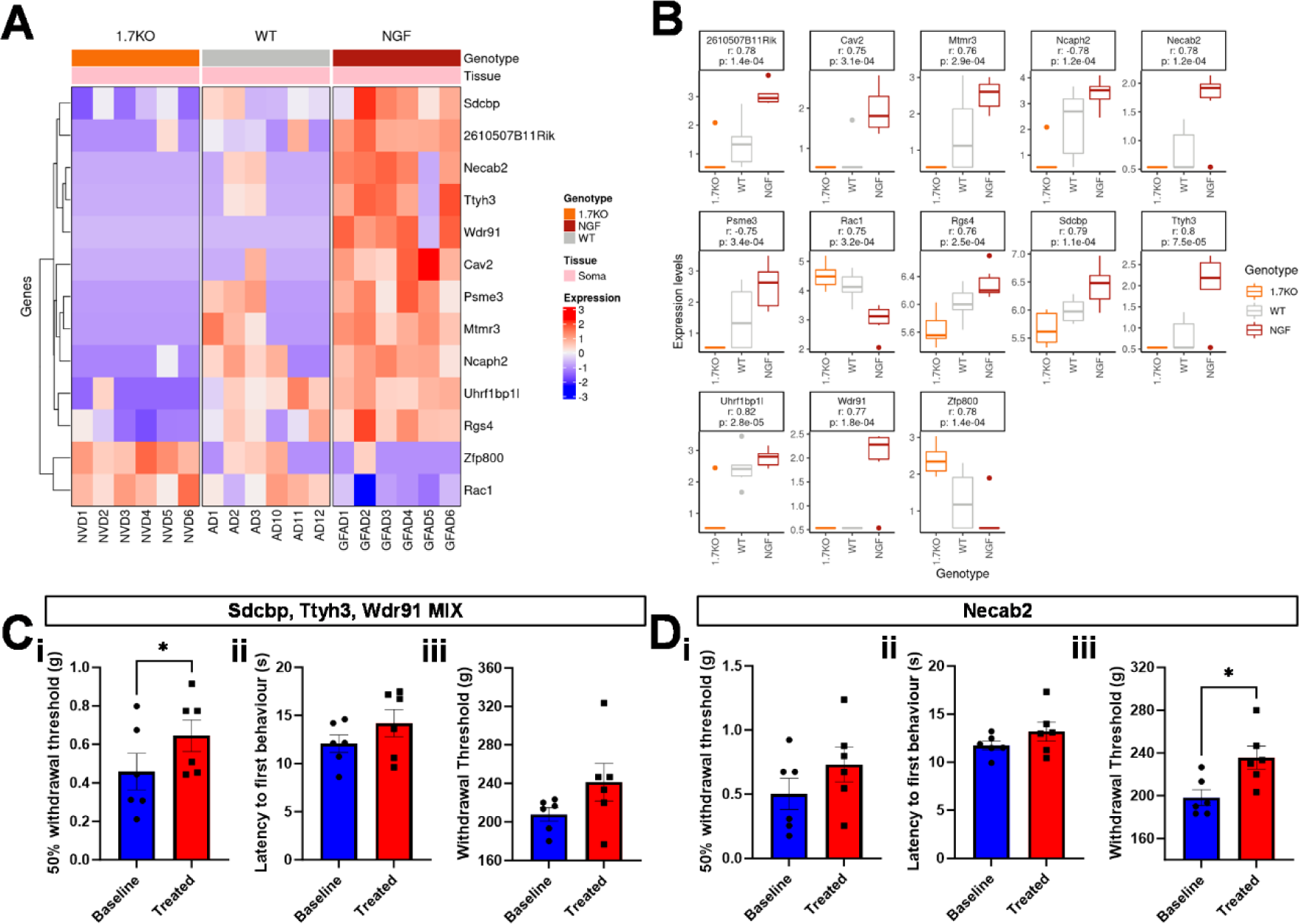
Pain-related somatic transcripts and effects of transcript ASO knockdown on acute pain behaviour. **A** Color-coded levels of expression in individual mice (abscissa) for transcripts that show a level of ribosome association with enhanced pain levels. **B** Relative levels of expression of transcripts shown in (**A**) derived from supplementary tables 4,6, and 8, together with Pearson’s correlation coefficient and the respective p value of the correlation. **C** ASO knockdown targeting combined Sdcbp, Ttyh3, and Wdr91 genes show a mild effect on mechanical sensation. **D** ASO knockdown targeting Necab2 leads to lowered acute mechanical pain behaviour. **i** = von Frey test, **ii** = Hargreaves’ heat test, **iii** = Randall Selitto test in both **C** and **D**. Motor impairment tests for **C** and **D** are shown in figure S5. Mean latencies and withdrawal thresholds in **C** and **D** were compared using paired t-test. * = p<0.05.

We decided to test an individual transcript *Necab2*, and in addition mix three sets of antisense oligonucleotides for a further experiment because we have a large number of potential targets, in order to minimise animal use. We selected *Sdcbp*, a syndecan-binding protein that shows excellent translational correlation with enriched expression after NGF treatment and lower expression in NaV1.7 null mice;^70^ *Ttyh3*,^52^ a large conductance calcium-activated chloride channel that presents high expression in NGF-treated mice and is absent in NaV1.7 nulls, and *Wdr91*, a negative regulator of the PI3 kinase activity which is selectively translated only after NGF-dependent pain induction. This mix showed a minor effect on mechanical thresholds using the Von Frey apparatus and Randall Selitto apparatus, and little to no effect on heat sensation (Figure 7C).

We also examined the effect of antisense oligonucleotides delivered intrathecally directed against the single target *Necab2*, which is NGF-induced and absent in pain-free samples (Figure 7D). This protein is a neuronal calcium-binding protein that binds to and modulates the function of two or more receptors, including adenosine A(2A) receptors and metabotropic glutamate receptor type 5 that are implicated in pain pathways. The inhibition of *Necab2* expression resulted in a diminution in noxious mechanosensation as measured with the Randall-Selitto apparatus, with little effect on heat sensing.

Changes in TRAP data from cell bodies contrasts with changes in central terminals, where proteins synthesised may be expected to play some role in interactions with spinal cord neurons and central pain pathways (Tables S8, S9). Once again, we exploited colour coded tables to summarise interesting transcripts that show pain-related expression. Two uncharacterised transcripts, *Gm4202* and *Gm10033*, showed an inverse correlation to pain and were not further investigated. *Zfp105* (ZNF35) is a retinoic acid regulated transcription factor.^71^ *Rps6ka3* is a member of the ribosomal 6 kinase family that play a wide-ranging role in signal transduction.^72^ BCR is a GTPase activating protein that also acts on tyrosine kinases. *Thoc7* has an important role in RNA translocation from the nucleus as well as viral release.^73^

We examined antisense knockdown of transcripts like *Thoc7* that show a pain correlation in terms of local protein synthesis (Figure 8).

**Figure 8:**
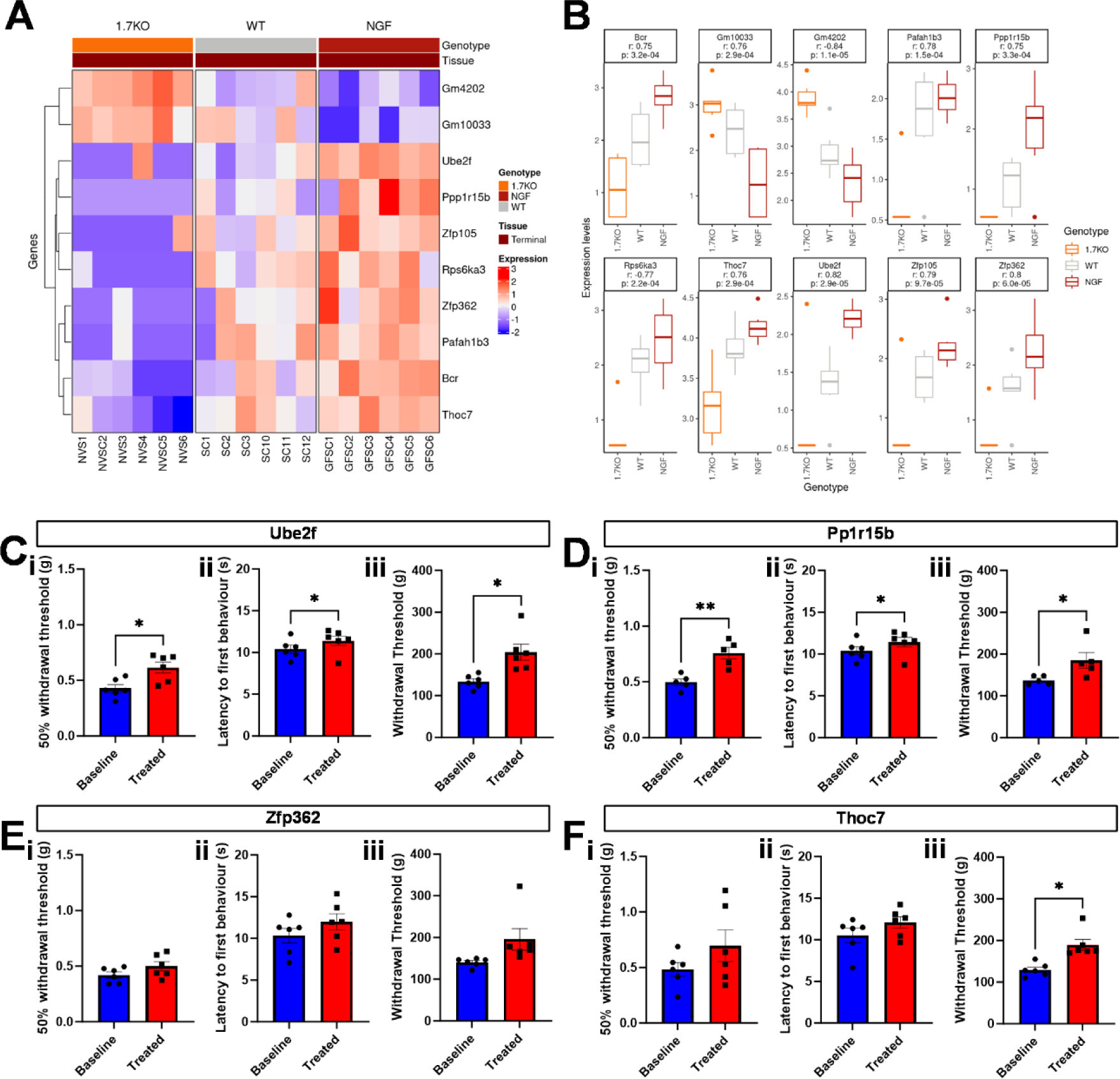
Pain-related central terminal transcripts and effects of transcript ASO knockdown on acute pain behaviour. **A** Color-coded levels of expression in individual mice (abscissa) for transcripts that show a level of ribosome association with enhanced pain levels. **B** Relative levels of expression of transcripts shown in (**A**) derived from supplementary tables 5, 7, and 9, together with Pearson’s correlation coefficient and the respective p value of the correlation. **C** ASO knockdown targeting Ube2f leads to lowered acute mechanical and thermal pain behaviour. **D** ASO knockdown targeting PP1r15b leads to lowered acute mechanical and thermal pain behaviour. **E** ASO knockdown targeting Zfp362 shows no significant change in acute pain behaviour. **F** ASO knockdown targeting Thoc7 leads to lowered acute mechanical pain behaviour. **i** = von Frey test, **ii** = Hargreaves’ heat test, **iii** = Randall Selitto test in **C**, **D**, **E**, and **F**. Motor impairment tests for **C**, **D**, **E**, and **F** are shown in figure S5. Mean latencies and withdrawal thresholds in **C**, **D**, **E** and **F** were compared using paired t-test. * = p<0.05; ** = p<0.01.

Other terminally translated proteins further investigated include *Ube2f*, a ubiquitin conjugating enzyme involved in neddylation that is known to play a role in pain pathways, and to stabilise voltage gated sodium channel activity. *Ppp1r15b* is a fascinating molecule that is regulated by glutamate receptor activity that controls its kinase or phosphatase inhibitory activity.^74^ Intriguingly, this protein can control the activity of *Eif2*, a key control for local translation. Thus, both proteins show a clear link to pain pathways. Lastly, *Zfp362* is assumed to be a transcription factor. *Pafah1b3* removes an acetyl group from PAF: its function is uncertain, but it seems to have a key role in brain development.^75^

Knockdown of both *Ube2f* and *Ppp1r15b* showed a significant reduction of both mechanical and thermal sensitivity across all the battery of tests used, highlighting even more their potential role in pain pathways (Figure 8C, D). *Thoc7* showed a significant increase in mechanical thresholds when using the Randall-Selitto apparatus, whereas knockdown of *Zfp362* did not show any effect on any of the pain tests performed (Figure 8E, F). For all antisense experiments, motor coordination was measured with a rotarod apparatus, and no impairment was observed (Figure S5).

## Discussion

Pain pathways depend on sensory neuron neurotransmitter input into the central nervous system evoked by damaging stimuli. The molecular organisation of the first synapse is incompletely understood and mainly rely on immunocytochemical studies. Protein synthesis is finely regulated at several stages, from transcription to translation and trafficking. Single cell RNA-seq has given helpful insights into cellular diversity and function, but there are limitations including the inability to detect low level transcripts, altered transcripts during circadian changes, and further changes caused by isolation of single cells from their normal milieu. Unfortunately, proteomic analysis is relatively insensitive, making single cell proteomic analysis at present problematic. There are thus significant gaps in our knowledge of the structure of the central terminals of sensory neurons, and potential changes that may occur in acute or chronic pain states. One interesting element in the physiology of sensory neurons is the existence of local protein synthesis at axon terminals. We have adapted the TRAP technology developed by Liu et al^18^ to examine proteins synthesised both in cell bodies and at the central terminals that may have a role in synaptic function. By comparing polysome-associated transcripts in pain states with those in normal or pain-free states, we have identified several mRNAs that could play a role in pain pathways. Interestingly, there is no obvious link between the translatome and the transcriptome in the somata of sensory neurons.^7^ Similarly, the mechanism and signals that result in mRNA translocation are poorly understood.^76^ There are no obvious consensus 5’ or 3’ UTR sequences linked to central terminal polysome-associated transcripts. We have used bioinformatic tools to classify genes that are co-ordinately regulated in response to altered pain states. These data are presented in Figure S6-S8.

A broad range of transcripts have been identified in this TRAP analysis, including data that suggest the vast majority of translatome transcripts are common to male and female mice. ncRNAs are present. These polysome-associated lncRNAs frequently have long ‘pseudo’ 5’UTRs and are 5’ capped, features that are important for ribosomal engagement and also nonsense-mediated decay.^37,77,78^ For the majority of known ribosome-associated lncRNAs, it is still unclear whether they are engaged by the ribosomes for translation (e.g. producing short/micro peptides),^79–81^ help to regulate protein translation,^82–85^ inertly reside in ribosomes or are degraded by the ribosome as a mechanism to control the cellular lncRNA population.^37^ However, for some specific lncRNAs their translation regulation function is known. For example, lincRNA-p21 associates with polysomes and suppresses the translation of JUNB and CTNNB1 mRNAs.^83^ In contrast, the natural antisense transcript to ubiquitin carboxy-terminal hydrolase L1 (AS-Uchl1), promotes the translation of Uchl1 by base pairing with its sense gene at a SINEB2 element and helps associate the sense mRNA with the active translating polysome.^82^

Genes that seem to play a role in pain pathways include a range of functional proteins. Mitochondrial genes that may play a role in the increased activity found in active sensory neurons signalling pain states have been identified (e.g.COX, see Figure S6). Other interesting transcripts that are clearly involved in pain pathways (e.g. *Penk*) show no altered levels of translatome association.^86^ The rate of translation of polysome associated genes may be regulated by kinase and phosphatases acting on eukaryotic initiation factors like EIF2, and such enzymes (e.g. *Ppp1r15b* are also present in the central translatome, and so relative read counts may not give a complete picture of effective translation.^61^

Bioinformatic analysis (Figure S8) shows that enhanced pain leads to more activity of genes involved in oxidative phosphorylation, synaptic vesicle activity, as one would suspect if sensory neurons were more actively signalling in pain states.

One striking observation is the presence of transcription factors in the central terminal translatome. *HoxC9* has been proposed to shuttle across membranes^87^ and other terminal Hox genes (for example *Hoxc8*) are implicated in motor neuron specification. It is an intriguing possibility that Hox proteins translated at sensory neuron terminals could exert functional effects on motor neurons.^88^

We have used antisense oligonucleotides to examine a role for some of the translatome mRNAs in acute pain and provide evidence that some do indeed contribute to pain states. These reagents may diminish mRNA levels in general rather than those associated with polysomes, but the net effect of diminishing protein production should be the same. We first checked two terminally enriched candidates, the G-protein regulatory protein *Gnas* and the Nitric Oxide synthase regulatory protein *Nos1ap*, both of which seem to play a role in acute pain. The novel genes that have so far not been linked to pain pathways were identified by correlating expression with enhanced pain states. Within the soma we found regulators of kainate receptor expression (*Sdcbp*) as well as metabotropic glutamate receptors (*Necab2*).^50,51^ A novel unique anion channel potentially activated by calcium (*Ttyh3*) was also identified as well as *Cav2*, a caveolin–like protein of unknown function. Both *Wdr91* and *Ncaph2* are implicated in neurodegeneration, whilst *Uhrf1bp1l* is a lipid transfer bridge and *Psme3* activates the proteosome. *Rgs4* regulates G-protein signalling and is a schizophrenia risk factor gene. *Rac1* is linked to innate immunity and the inflammasome whilst *Zfp362* is a potential transcription factor without activity in acute pain. Tables S4-S9 contain complete rank ordered lists of altered genes with P values and read counts.

Our functional studies demonstrate a clear role in acute pain for the terminally enriched transcripts *Gnas* and *Nos1Ap*, as well as *Ppp1r15b* that regulates translational activity via Eif proteins, *Thoc7* that regulates RNA transport and *Ube2f* a regulator of neddylation are also linked to pain. Neither scrambled ASO controls nor *Zfp362* ASOs showed any effect on pain behaviour (Figure 8E, Figure S3). Within the soma, *Necab2* was a strong candidate that showed a major effect on mechanical pain. With the current availability of targeted delivery systems and advances in gene therapy, these targets are worthy of further study in other models of human chronic pain. In the future it would be helpful to appraise more complex pain states like chronic inflammatory and neuropathic pain with the substantial number of candidate genes identified, but in the interest of minimising animal suffering, candidates that have some genetic links to human pain (e.g. *Ttyh3*) should be prioritised.

## Acknowledgements

We acknowledge with gratitude the following sources of funding: Versus Arthritis UK (21950), Medical Research Council (MR/V012509/1; 571476), and Cancer Research UK (185341). This work acknowledges the support of the National Institute for Health Research Barts Biomedical Research Centre (NIHR 203330). The views expressed are those of the authors and do not represent those of the NHS, NIHR or funding bodies.

## Author Contributions

Conceptualization, M.A.B., J.Z., and J.N.W.; Formal Analysis, C.C., M.J.L.; Investigation, M.A.B., F.I., A.P.L., M.A., R.H., S.J.G., S.S-V.; Resources, F.I., R.H., S.J.G., S.S-V; Data Curation, C.C.; Visualization, F.I., C.C., M.A.B.; Writing – Original Draft, J.N.W., J.Z., C.C., F.I.; Writing – Review and Editing, F.I., R.H., J.J.C., M.J.L., J.Z., J.N.W.; Supervision, J.N.W., J.Z., M.J.L.; Funding Acquisition, J.N.W., M.J.L.;

## STAR METHODS

### KEY RESOURCES TABLE

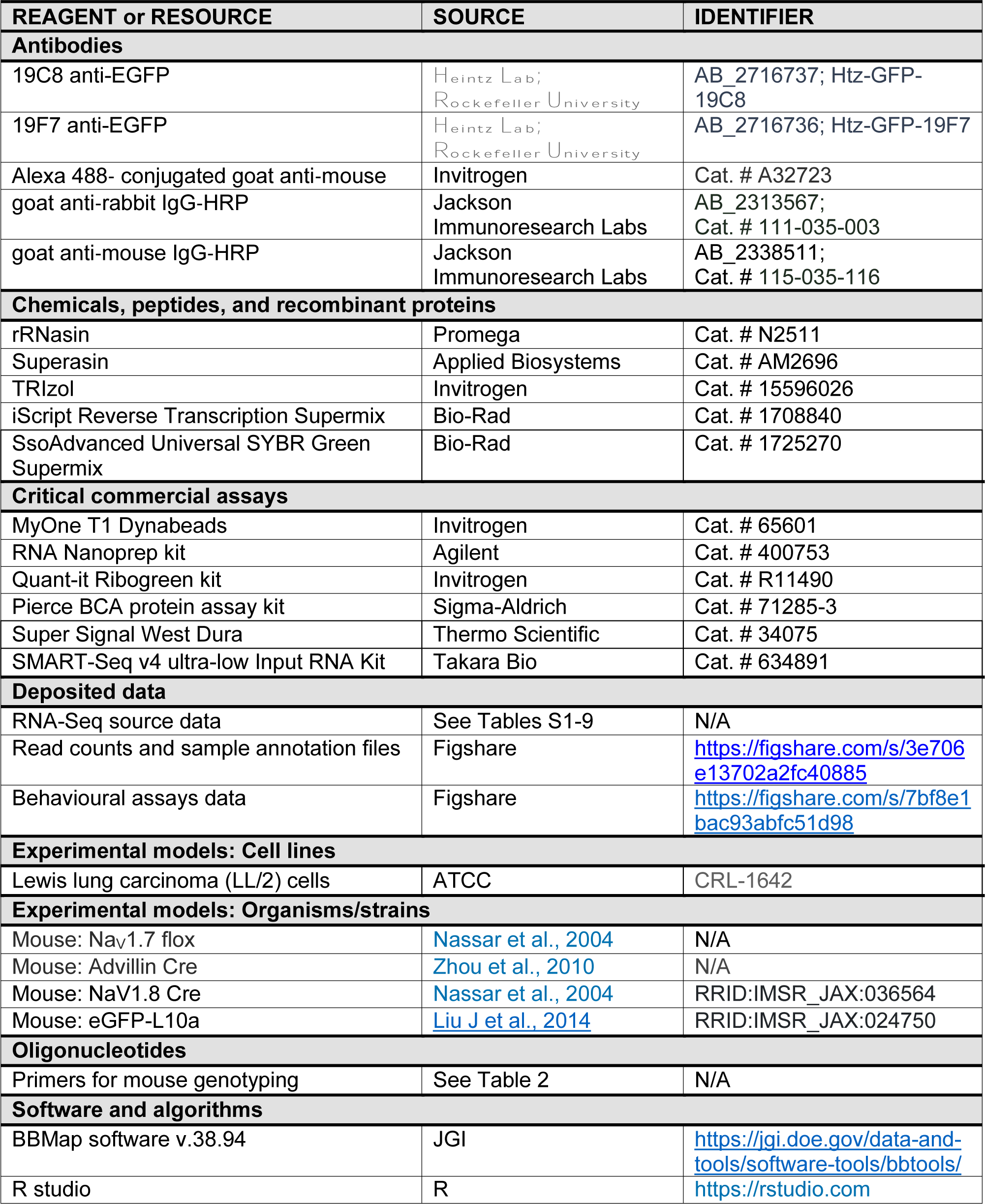
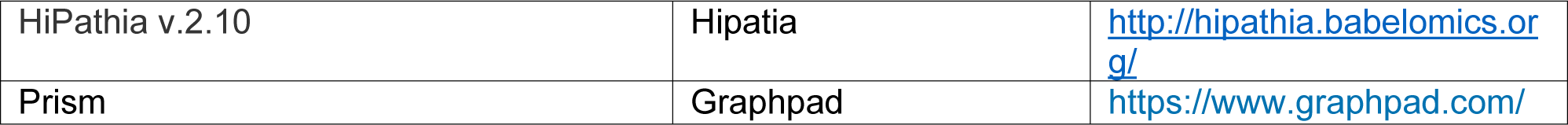

## RESOURCE AVAILABILITY

### Lead contact

Further information and requests for resources and reagents should be directed to and will be fulfilled by the lead contact, John N. Wood.

### Materials availability

This study did not generate any new unique reagents.

### Data and code availability

Data is available from the lead contact on reasonable request.

Complete data comprising read numbers, fold increase (log_2_) and p-values are presented in tables S1-S9.

TRAP gene expression counts and sample annotation files, as well as behavioural assays data files, are deposited on Mendeley data at https://data.mendeley.com/preview/ppj2ptnpgz?a=0b4c72a3-e21c-4bf3-a8e4-1d0e5b7abcc9.

## EXPERIMENTAL MODEL DETAILS

### Animals

All experiments were performed in compliance with the UK Animals (Scientific Procedures) Act 1986, under a Home Office project licence (PPL 413329A2). Mice were housed in a 12-h light/dark cycle with *ad libitum* provision of food and water. All mice were acclimatized for 1 week to the facility before the start of experiments. Mice were housed in individually ventilated cages (Techniplast GM500 Mouse IVC Green line) containing Lignocel bedding with a maximum of 5 adult mice per cage. All experiments were carried out using adult male and female mice. Mice were euthanized by CO_2_ asphyxiation followed by cervical dislocation.

Mice in pain free and painful conditions were tested, using Nav1.7 null mutant mice generated by crossing floxed Nav1.7 with Advillin-Cre. Mice in pain were generated by systemic injection of NGF.

### Generation of eGFP-L10a mice

eGFP-L10a mice were obtained from the Jackson laboratory. Homozygous or Heterozygous eGFP-L10a were then crossed with either Nav1.8-cre or Adv-cre mice to obtain Cre; eGFP-L10a heterozygote mice. All experiments were performed in 8-12 weeks old male and female mice. Details of primers used for genotyping are available in table 1.

**Table 1:**
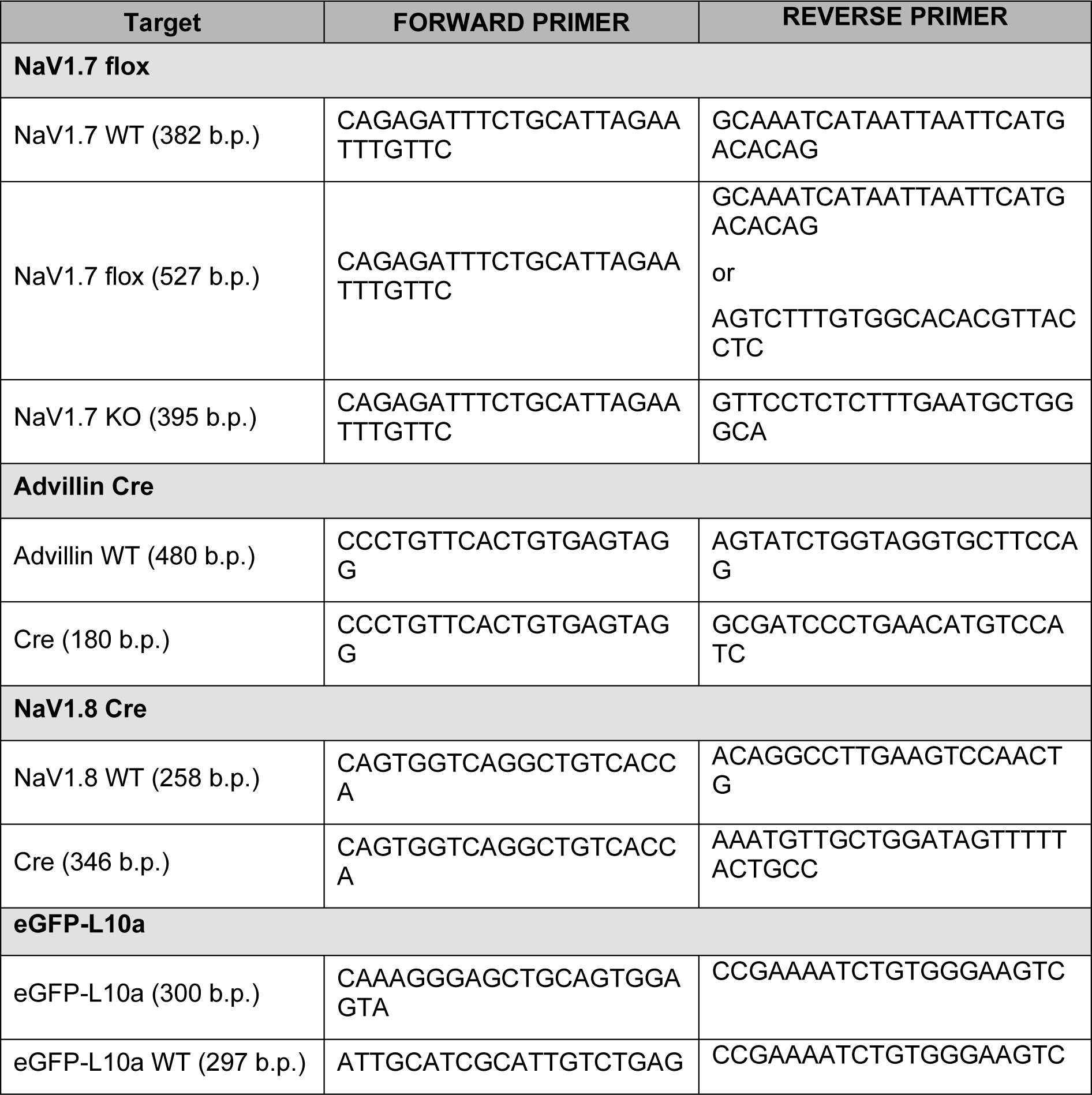
Primers used for genotyping. Sequences are listed as 5’-3’.

## METHOD DETAILS

### Translating Ribosome Affinity Purification (TRAP)

TRAP assay was performed as described^89^ with slight modifications. Briefly, affinity matrix was prepared by incubating MyOne T1 Dynabeads (Life Technology) with biotinylated protein L and monoclonal 19C8 and 19F7 eGFP antibodies^90^. All the bilateral DRGs or Spinal Cords from 3 male or female mice were dissected on ice, pooled and homogenized in low salt buffer, followed by extraction of post-nuclear fraction at 1000 *g* spin. This fraction was then homogenized in non-denaturing 1% NP-40 buffer followed by isolation of post-mitochondrial fraction at 12000 *g* spin. GFP tagged ribosomes were then isolated by mixing the post-mitochondrial fraction with the pre-prepared affinity matrix of eGFP antibodies and beads overnight at 4°C. After washing the beads in high-salt buffer, the ribosome bound RNA was eluted and purified using RNA Nanoprep kit (Agilent). Finally, the isolated ribosomal bound RNA was quantified using Quant-it Ribogreen kit (Invitrogen). All buffers had 100 ug/ml cycloheximide (Sigma) with 10ul/ml rRNasin (Promega) and Superasin (Applied Biosystems) to inhibit RNAases, while all reactions were carried out in RNAse-free tubes (Ambion).

### Reverse Transcriptase (RT)-qPCR

DRGs from all the segments or spinal cords were dissected from three mice and pooled. RNA was extracted using TRIzol Reagent (Invitrogen) according to the manufacturer’s instructions. Reverse transcription was performed using iScript Reverse Transcription Supermix for reverse transcriptase–qPCR following the supplied protocol by Invitrogen. Complementary DNA amplification was performed in triplicate, using SsoAdvanced Universal SYBR Green Supermix (Bio-Rad).

### Immunohistochemistry

Following anaesthesia, mice were trans-cardially perfused with PBS, followed by 4% paraformaldehyde in PBS. DRGs and spinal cord were dissected and incubated in fixative for 4h at 4°C, followed by 30% sucrose in PBS overnight at 4°C. Tissue was embedded in O.C.T. (Tissue-Tek) and snap-frozen in a dry ice/2-methylbutane bath. DRG and spinal cord cross cryosections (20μm) were collected on glass slides (Superfrost Plus, Polyscience) and stored at −80°C until further processing. For GFP-tag immunohistochemistry in DRGs and spinal cord, sections were incubated in blocking solution (4% horse serum, 0.3% Triton X-100 in PBS) for 1h at room temperature, followed by incubation in mouse anti-GFP antibody (C7 or F8) diluted 1:200 in blocking solution overnight at 4°C. Following three washes in PBS, bound antibody was visualised using an Alexa 488-conjugated goat anti-mouse secondary antibody (1:1000, Invitrogen). Fluorescence images were acquired on confocal laser scanning microscope (LSM 780, Zeiss).

### Immunoblotting

Proteins for immunoblots were isolated from freshly excised DRG or spinal cord followed by homogenization in RIPA lysis buffer as described previously^91^. The nuclear fraction and cell debris were removed by centrifugation at ∼20,000 *g* for 15 min at 4°C. Protein concentrations were determined with Pierce BCA protein assay kit, and then samples of 40 μg were separated on SDS–PAGE gel in Bio-Rad Mini-PROTEAN Vertical Electrophoresis Cell System and blotted to the Immobilin-P membrane (IPVH00010, Millipore) in transfer buffer (25 mM Tris–HCl, pH 8.3, 192 mM glycine, 0.1% SDS and 20% methanol) for 1 h at 100 V with a Bio-Rad transfer cell system. The membrane was blocked in blocking buffer [5% nonfat milk in PBS–Tween buffer (0.1% Tween 20 in 1× PBS)] for 1 h at room temperature and then incubated with primary antibody, anti-GFP (C7, 1:1000) in blocking buffer overnight at 4°C. The membrane was washed three times with TBS–Tween (20 mM Tris, 150 mM NaCl, 0.1% Tween 20, pH 7.5) and then incubated with secondary antibody goat anti-mouse or goat anti-rabbit IgG-HRP (1:4,000; Jackson ImmunoResearch Laboratories) in TBS–Tween at room temperature for 2 h. Detection was performed using a Western Lightning Chemiluminescence Reagent (Super Signal Western Dura, Thermo Scientific) and exposed to BioMax film (Kodak).

### RNA-Seq Analysis

RNA integrity of ribosome-bound mRNA samples from three replicates was assessed using RIN and DV200 scores. SMART-Seq v4 ultra-low RNA (Takara Bio) library preparation protocol was used to generate cDNA libraries and sequencing was performed on an Illumina HiSeq 2 instrument with 150 bp reads according to the manufacturer’s instructions. Samples with 50M reads were sequenced using a 2×150-base-pair (BP) paired-end configuration. After demultiplexing, Illumina adapters and nucleotides with poor quality were trimmed using bbduk from the BBMap software v.38.94. The mouse reference genome, Gencode release v28 (GRCm39), was edited by concatenation of the eGFP sequence retrieved from https://www.ncbi.nlm.nih.gov/nuccore/EU056363.1. The reference genome indexing, read mapping and counting mapped reads were performed using the STAR aligner v.2.7.3a in the 2-pass mapping mode that allows for unbiased exon splice junction detection.

After extracting uniquely mapped read counts, the genes that showed low expression across all samples, less than 20 total counts, were discarded from our dataset and the remaining genes were used for data quality control and downstream analysis. The remaining counts were subjected to variance-stabilizing transformation (VST) and then differential expression analysis was performed between different conditions using the DESeq2 v.1.25.9 package. A Principal Component Analysis (PCA) for unsupervised projection of RNA-Seq data was performed with prcomp function in R using normalized data and PCA plots were generated using ggplot2 v.3.3.6. All the sample clusters shown in PCA plots were highly associated with biological conditions rather than technical ones. Therefore, no batch effect correction was applied. Partition of samples based on the proportion of neuronal mRNA content was assessed using 32 genes recently published by Pradipta R Ray et al.^92^. The median value of these genes was used as neuronal module score and then samples were categorized into three groups (enriched, moderate and de-enriched) using the 30th and 70th percentiles of module scores. DESeq2 was used to calculate p-values (statistical analysis by negative binomial general linear model with Wald test) and log_2_ fold changes that were used to generate volcano plots using easylabel package v.0.2.4. Linear relationships between genes and three genotypes (1.7KO, WT, NGF / −1, 0, 1) were identified by means of Pearson’s correlation using the cor.test function of the R Stats package. Results are shown for only the genes with absolute correlation coefficient > 0.75 and p-value < 0.05. Heatmaps were generated using ComplexHeatmap v.2.12.1.

### Pathway Analysis

#### Geneset Enrichment Analysis

Differentially expressed genes were used for gene set enrichment analysis. This analysis was performed with an R interface to the Enrichr database v.3.0. The hypergeometric model was used to assess whether the number of selected genes associated with a KEGG pathway was larger than expected. Pathway terms with p-value < 0.05 were considered significant.

#### Differentially Active Pathways

The expression levels of the genes corresponding to the proteins involved in the pathways are used by the mechanistic pathway models to infer the activities of the pathways. Activities of signaling and metabolic sub-pathways were estimated using Hipathia v.2.10 and Metabolizer v.1.7.0 tools, respectively. A two-sided Wilcoxon signed-rank test was used for the statistical assessment when comparing different conditions.

#### Antisense Oligonucleotides

Antisense Oligonucleotides (ASOs) directed against sequences encompassing initiator methionines and polyadenylation sites of candidate mRNAs were purchased from Sigma-Aldrich. All ASOs were 20mers, HPLC purified with phosphorothioate nucleotides at the 5′ and 3′ end (see Table 2)^93^.

**Table 2:**
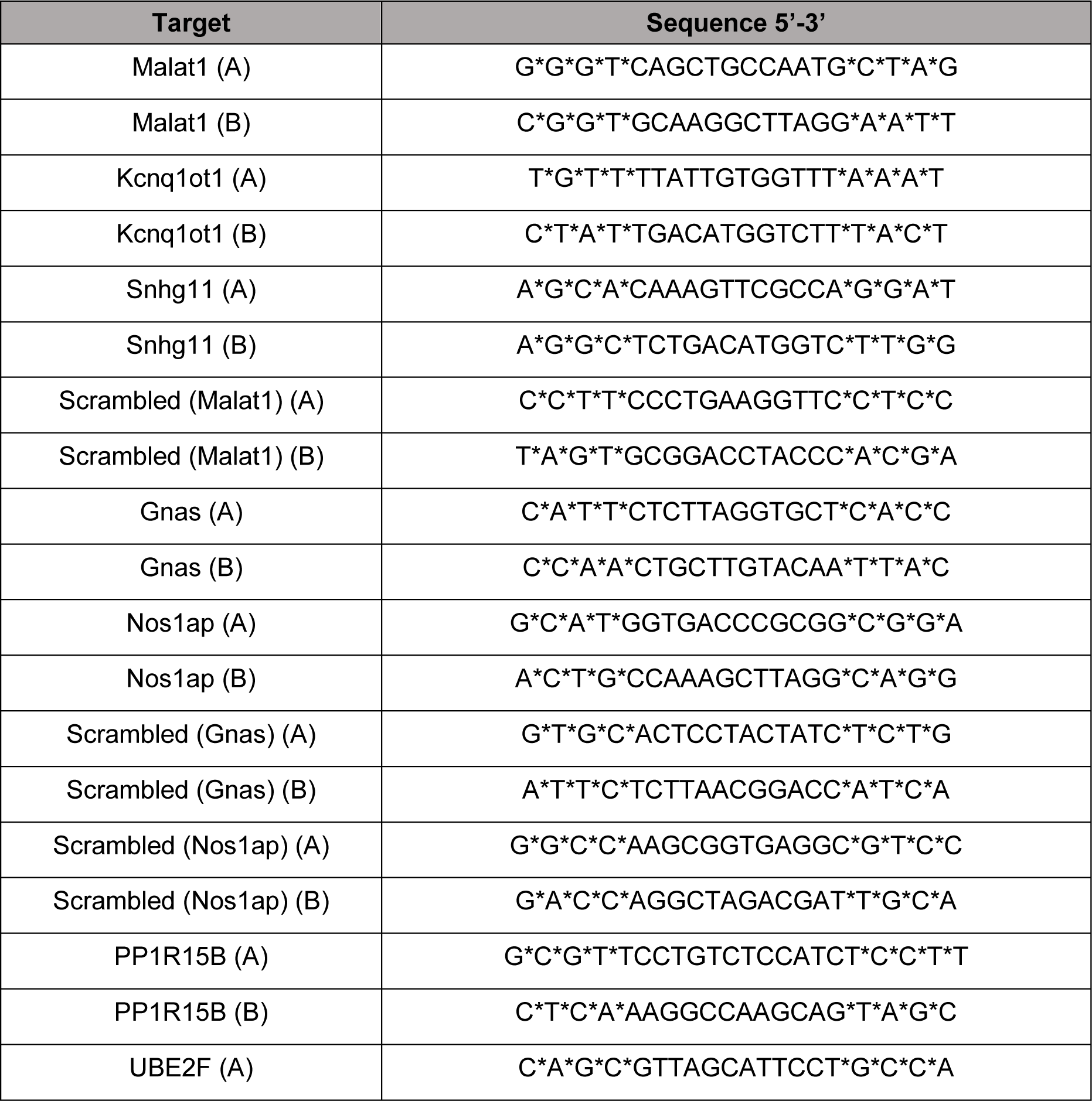
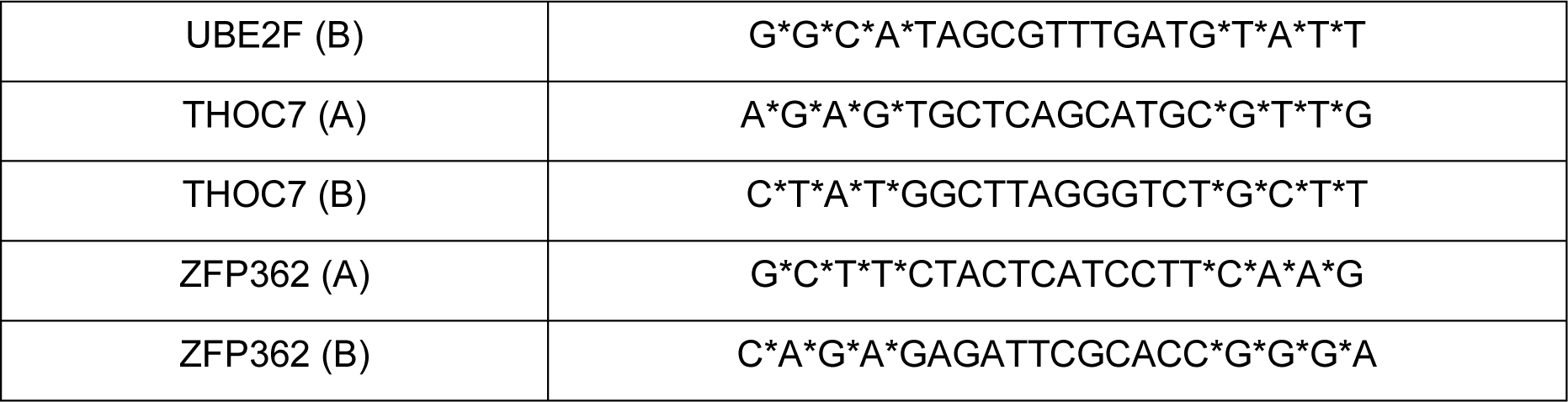
Sequence of each pair of targeted and control ASO used. Asterisk denotes phosphorothioate residues; target species is mouse.

#### ASOs Injections

Adult C57BL/6J mice between 8-10 weeks were anaesthetised using 2-3% Isoflurane and injected with 6 µl of two or more targeted of control antisense oligonucleotides resuspended in PBS for a total amount of 15 µg/mouse/injection via the intrathecal route using a Hamilton syringe connected to a 30G needle cannula. A pair of ASOs designed for the extremities of the mRNA were designed for each target: their sequences can be found in Table 2. The ASOs were injected on 3 days (every other day), and mice were prepared for behavioural testing immediately after the last injection.

#### Behavioural Testing

All animal experiments were performed in accordance with Home Office Regulations. Observers were blinded to treatment. Animals were acclimatized to handling by the investigator and every effort was made to minimize stress during the testing. Male animals were used for experiments apart from gender comparison studies. All experiments were filmed, and carried out by two independent researchers^94^. Cancer pain studies were carried out exactly as described in ^95^.

#### Randall Selitto

The threshold for mechanonociception was assessed using the Randall Selitto test^96^ Animals were restrained in a clear plastic tube. A 3 mm^2^ blunt probe was applied to the tail of the animal with increasing pressure until the mouse exhibited a nocifensive response, such as tail withdrawal. The pressure required to elicit the nocifensive behavior was averaged across three trials. The cut-off was set to 500 g.

#### Von Frey

Punctate mechanical sensitivity was measured using the up-down method of Chaplan to obtain a 50% withdrawal threshold^97^. Mice were habituated for one hour in darkened enclosures with a wire mesh floor. A 0.4 g von Frey filament was applied to the plantar surface of the paw for 3 s. A positive response resulted in application of a filament of lesser strength on the following trial, and no response in application of a stronger filament. To calculate the 50% withdrawal threshold, five responses surrounding the 50% threshold were obtained after the first change in response. The pattern of responses was used to calculate the 50% threshold = (10[χ+κδ])/10,000), where χ is the log of the final von Frey filament used, κ = tabular value for the pattern of responses and δ the mean difference between filaments used in log units. The log of the 50% threshold was used to calculate summary and test statistics, in accordance with Weber’s Law.

#### Hargreaves’ Test

Spinal reflex responses to noxious heat stimulation were assessed using the Hargreaves’ test^98^. Mice were habituated for an hour in plexiglass enclosures with a glass base. Before testing, the enclosures were cleaned of faeces and urine. Radiant heat was then locally applied to the plantar surface of the hindpaw until the animal exhibited a nocifensive withdrawal response. Average latencies were obtained from three trials per animal, with inter-trial interval of 15 mins. Cut-off time was set to 30 s.

### Cancer-Induced Bone Pain

#### Cell culture

Lewis lung carcinoma (LL/2) cells (from American Type Culture Collection (ATCC)) were cultured in a medium containing 90% Dulbecco’s Modified Eagle Medium (DMEM) and 10% fetal bovine serum (FBS) and 0.1% Penicillin/Streptomycin for 14 days before the surgery. DMEM was supplemented with L-glutamine (1%) and glucose (4.5 g/l). The cells were sub-cultured whenever ∼80% confluency was reached, which was done a day before the surgery. On the surgery day, LL/2 cells were harvested by scraping and were centrifuged at a speed of 1500 rpm for two mins. The supernatant was removed, and the cells were resuspended in a culture medium that contained DMEM to attain a final concentration of ∼2×10^6^ cells/ml. The cell counting and viability check were done using the Countess automated cell counter (Thermo Fisher Scientific).

#### Surgery

Surgery was carried out on anaesthetized C57BL/6 mice. Anesthesia was achieved using 2-3% isoflurane. The legs and the thighs of the mice were shaved, and the shaved area was cleaned using hibiscrub solution. A sterile lacri-lube was applied to the eyes and lidocaine was applied at the site of the surgery. The reflexes of the mice to pinches were checked to ensure successful anaesthesia. An incision was made in the skin above and lateral to the patella on the left leg. The patella and the lateral retinaculum tendons were loosened to move the patella to the side and expose the distal femoral epiphysis. A 30G needle was used to drill a hole through the femur to permit access to the intramedullary space of the femur. The 30G needle was removed, and a 0.3ml insulin syringe was used to inoculate ∼2×10^4^ LL/2 cells suspended in DMEM. The hole in the distal femur was sealed using bone wax (Johnson & Johnson). To ensure that there was no bleeding, the wound was washed with sterile normal saline. Following that, the patella was placed back into its original location, and the skin was sutured using 6–0 absorbable vicryl rapid (Ethicon). Lidocaine was applied at the surgery site, and the animals were placed in the recovery chamber and monitored until they recovered.

#### Limb-use score

The mice housed in the same cage were placed together in a glass box (30 × 45 cm) for at least five minutes. Then each mouse was left in the glass box on its own and was observed for ∼4 minutes, and the use of the ipsilateral limb was estimated using the standard limb use scoring system in which: 4 indicates a normal use of the affected limb; 3 denotes slight limping (slight preferential use of the contralateral limb when rearing); 2 indicates clear limping; 1 clear limping, and with a tendency of not using the affected limb; and 0 means there is no use of the affected limb. Reaching a limb-use score of zero was used as an endpoint for the experiment.

#### Static weight-bearing

The scale used for this behavioural test was the Incapacitance Metre (Linton Instrumentation), which has two scales to assess the weight put on each limb. The weight placed on each limb is measured for 3 seconds. The readings for the weight put on the limbs were recorded three times for each mouse, and between the readings, mice were allowed to re-place themselves into the tube. The fraction of the weight put on the ipsilateral paw was determined by the summation of all three readings of the weight put on the ipsilateral paw divided by the summation of all the weight measurements on both paws.

## QUANTIFICATION AND STATISTICAL ANALYSIS

For behavioural experiments, *n* refers to the number of animals.

Details about the statistical and correlation analyses of RNA-Seq data are detailed in the appropriate methods sections.

Datasets are presented using appropriate summary statistics as indicated in the figure legends. Error bars in all graphs denote mean ± SEM. Tests of statistical comparison for each dataset are described in detail in the figure legends. For grouped data, we made the appropriate correction for multiple comparisons. We set an α-value of p=0.05 for significance testing and report all p values resulting from planned hypothesis testing.

## References

1. Akopian, A.N., and Wood, J.N. (1995). Peripheral nervous system-specific genes identified by subtractive cDNA cloning. J Biol Chem 270, 21264–21270.

2. Bangash, M.A., Alles, S.R.A., Santana-Varela, S., Millet, Q., Sikandar, S., de Clauser, L., Ter Heegde, F., Habib, A.M., Pereira, V., Sexton, J.E., et al. (2018). Distinct transcriptional responses of mouse sensory neurons in models of human chronic pain conditions. Wellcome Open Res 3, 78. 10.12688/wellcomeopenres.14641.1.

3. LaCroix-Fralish, M.L., Austin, J.S., Zheng, F.Y., Levitin, D.J., and Mogil, J.S. (2011). Patterns of pain: meta-analysis of microarray studies of pain. Pain 152, 1888–1898. 10.1016/j.pain.2011.04.014.

4. Kanellopoulos, A.H., Koenig, J., Huang, H., Pyrski, M., Millet, Q., Lolignier, S., Morohashi, T., Gossage, S.J., Jay, M., and Linley, J.E. (2018). Mapping protein interactions of sodium channel NaV1. 7 using epitope-tagged gene-targeted mice. The EMBO journal 37, 427-445.

5. Huang, H.-L., Cendan, C.-M., Roza, C., Okuse, K., Cramer, R., Timms, J.F., and Wood, J.N. (2008). Proteomic profiling of neuromas reveals alterations in protein composition and local protein synthesis in hyper-excitable nerves. Molecular pain 4, 1744–8069-1744-1733.

6. Usoskin, D., Furlan, A., Islam, S., Abdo, H., Lonnerberg, P., Lou, D., Hjerling-Leffler, J., Haeggstrom, J., Kharchenko, O., Kharchenko, P.V., et al. (2015). Unbiased classification of sensory neuron types by large-scale single-cell RNA sequencing. Nat Neurosci 18, 145–153. 10.1038/nn.3881.

7. Schmidt, M., Sondermann, J.R., Gomez-Varela, D., Çubuk, C., Millet, Q., Lewis, M.J., Wood, J.N., and Zhao, J. (2022). Transcriptomic and proteomic profiling of NaV1. 8-expressing mouse nociceptors. Frontiers in Molecular Neuroscience 15, 1002842.

8. Shigeoka, T., Koppers, M., Wong, H.H., Lin, J.Q., Cagnetta, R., Dwivedy, A., de Freitas Nascimento, J., van Tartwijk, F.W., Strohl, F., Cioni, J.M., et al. (2019). On-Site Ribosome Remodeling by Locally Synthesized Ribosomal Proteins in Axons. Cell Rep 29, 3605–3619 e3610. 10.1016/j.celrep.2019.11.025.

9. Scarnati, M.S., Kataria, R., Biswas, M., and Paradiso, K.G. (2018). Active presynaptic ribosomes in the mammalian brain, and altered transmitter release after protein synthesis inhibition. Elife 7. 10.7554/eLife.36697.

10. Sahoo, P.K., Lee, S.J., Jaiswal, P.B., Alber, S., Kar, A.N., Miller-Randolph, S., Taylor, E.E., Smith, T., Singh, B., and Ho, T.S.-Y. (2018). Axonal G3BP1 stress granule protein limits axonal mRNA translation and nerve regeneration. Nature communications 9, 3358.

11. MacDonald, D.I., Sikandar, S., Weiss, J., Pyrski, M., Luiz, A.P., Millet, Q., Emery, E.C., Mancini, F., Iannetti, G.D., and Alles, S.R. (2020). The mechanism of analgesia in NaV1. 7 null mutants. bioRxiv, 2020.2006. 2001.127183.

12. MacDonald, D.I., Sikandar, S., Weiss, J., Pyrski, M., Luiz, A.P., Millet, Q., Emery, E.C., Mancini, F., Iannetti, G.D., and Alles, S.R. (2021). A central mechanism of analgesia in mice and humans lacking the sodium channel NaV1. 7. Neuron 109, 1497-1512.e1496.

13. Heiman, M., Schaefer, A., Gong, S., Peterson, J.D., Day, M., Ramsey, K.E., Suarez-Farinas, M., Schwarz, C., Stephan, D.A., and Surmeier, D.J. (2008). A translational profiling approach for the molecular characterization of CNS cell types. Cell 135, 738–748.

14. Doyle, J.P., Dougherty, J.D., Heiman, M., Schmidt, E.F., Stevens, T.R., Ma, G., Bupp, S., Shrestha, P., Shah, R.D., and Doughty, M.L. (2008). Application of a translational profiling approach for the comparative analysis of CNS cell types. Cell 135, 749–762.

15. Rozenbaum, M., Rajman, M., Rishal, I., Koppel, I., Koley, S., Medzihradszky, K.F., Oses-Prieto, J.A., Kawaguchi, R., Amieux, P.S., Burlingame, A.L., et al. (2018). Translatome Regulation in Neuronal Injury and Axon Regrowth. eNeuro 5. 10.1523/ENEURO.0276-17.2018.

16. Megat, S., Ray, P.R., Moy, J.K., Lou, T.F., Barragan-Iglesias, P., Li, Y., Pradhan, G., Wanghzou, A., Ahmad, A., Burton, M.D., et al. (2019). Nociceptor Translational Profiling Reveals the Ragulator-Rag GTPase Complex as a Critical Generator of Neuropathic Pain. J Neurosci 39, 393–411. 10.1523/JNEUROSCI.2661-18.2018.

17. Tavares-Ferreira, D., Ray, P.R., Sankaranarayanan, I., Mejia, G.L., Wangzhou, A., Shiers, S., Uttarkar, R., Megat, S., Barragan-Iglesias, P., Dussor, G., et al. (2020). Sex Differences in Nociceptor Translatomes Contribute to Divergent Prostaglandin Signaling in Male and Female Mice. Biol Psychiatry. 10.1016/j.biopsych.2020.09.022.

18. Liu, J., Krautzberger, A.M., Sui, S.H., Hofmann, O.M., Chen, Y., Baetscher, M., Grgic, I., Kumar, S., Humphreys, B., and Hide, W.A. (2014). Cell-specific translational profiling in acute kidney injury. The Journal of clinical investigation 124, 1242–1254.

19. Santana-Varela, S., Bogdanov, Y.D., Gossage, S.J., Okorokov, A.L., Li, S., De Clauser, L., Alves-Simoes, M., Sexton, J.E., Iseppon, F., and Luiz, A.P. (2021). Tools for analysis and conditional deletion of subsets of sensory neurons. Wellcome Open Research 6.

20. Barker, P.A., Mantyh, P., Arendt-Nielsen, L., Viktrup, L., and Tive, L. (2020). Nerve growth factor signaling and its contribution to pain. Journal of pain research, 1223-1241.

21. Nguyen, E., Lim, G., Ding, H., Hachisuka, J., Ko, M.-C., and Ross, S.E. (2021). Morphine acts on spinal dynorphin neurons to cause itch through disinhibition. Science Translational Medicine 13, eabc3774.

22. Megat, S., Ray, P.R., Moy, J.K., Lou, T.-F., Barragán-Iglesias, P., Li, Y., Pradhan, G., Wanghzou, A., Ahmad, A., and Burton, M.D. (2019). Nociceptor translational profiling reveals the Ragulator-Rag GTPase complex as a critical generator of neuropathic pain. Journal of Neuroscience 39, 393–411.

23. Rozenbaum, M., Rajman, M., Rishal, I., Koppel, I., Koley, S., Medzihradszky, K.F., Oses-Prieto, J.A., Kawaguchi, R., Amieux, P.S., and Burlingame, A.L. (2018). Translatome regulation in neuronal injury and axon regrowth. Eneuro 5.

24. Tavares-Ferreira, D., Ray, P.R., Sankaranarayanan, I., Mejia, G.L., Wangzhou, A., Shiers, S., Uttarkar, R., Megat, S., Barragan-Iglesias, P., and Dussor, G. (2022). Sex differences in nociceptor translatomes contribute to divergent prostaglandin signaling in male and female mice. Biological Psychiatry 91, 129–140.

25. Schwämmle, T., and Schulz, E.G. (2023). Regulatory principles and mechanisms governing the onset of random X-chromosome inactivation. Current Opinion in Genetics & Development 81, 102063.

26. Brooks, S.P., Coccia, M., Tang, H.R., Kanuga, N., Machesky, L.M., Bailly, M., Cheetham, M.E., and Hardcastle, A.J. (2010). The Nance–Horan syndrome protein encodes a functional WAVE homology domain (WHD) and is important for co-ordinating actin remodelling and maintaining cell morphology. Human molecular genetics 19, 2421–2432.

27. Zhou, Z.-D., Saw, W.-T., and Tan, E.-K. (2017). Mitochondrial CHCHD-containing proteins: physiologic functions and link with neurodegenerative diseases. Molecular neurobiology 54, 5534–5546.

28. Zinn, A.R., Alagappan, R.K., Brown, L.G., Wool, I., and Page, D.C. (1994). Structure and function of ribosomal protein S4 genes on the human and mouse sex chromosomes. Molecular and cellular biology.

29. Zhang, M., Zhou, Y., Jiang, Y., Lu, Z., Xiao, X., Ning, J., Sun, H., Zhang, X., Luo, H., and Can, D. (2021). Profiling of sexually dimorphic genes in neural cells to identify Eif2s3y, whose overexpression causes autism-like behaviors in male mice. Frontiers in Cell and Developmental Biology 9, 669798.

30. Soto, F., Tien, N.-W., Goel, A., Zhao, L., Ruzycki, P.A., and Kerschensteiner, D. (2019). AMIGO2 scales dendrite arbors in the retina. Cell reports 29, 1568–1578. e1564.

31. Goodman, E., and Iversen, L. (1986). Calcitonin gene-related peptide: novel neuropeptide. Life sciences 38, 2169–2178.

32. Lischka, A., Eggermann, K., Record, C.J., Dohrn, M.F., Laššuthová, P., Kraft, F., Begemann, M., Dey, D., Eggermann, T., and Beijer, D. (2023). Genetic landscape of congenital insensitivity to pain and hereditary sensory and autonomic neuropathies. Brain, awad328.

33. Grone, B., and Maruska, K. (2016). Three distinct glutamate decarboxylase genes in vertebrates. Sci Rep 6: 30507.

34. Lee, W.-H., Li, L.-L., Chawla, A., Hudmon, A., Lai, Y.Y., Courtney, M.J., and Hohmann, A.G. (2018). Disruption of nNOS-NOS1AP protein-protein interactions suppresses neuropathic pain in mice. Pain 159, 849.

35. Zhadina, M., Roszko, K.L., Geels, R.E., de Castro, L.F., Collins, M.T., and Boyce, A.M. (2021). Genotype-phenotype correlation in fibrous dysplasia/McCune-Albright syndrome. The Journal of Clinical Endocrinology & Metabolism 106, 1482–1490.

36. Carlevaro-Fita, J., Rahim, A., Guigó, R., Vardy, L.A., and Johnson, R. (2016). Cytoplasmic long noncoding RNAs are frequently bound to and degraded at ribosomes in human cells. Rna 22, 867–882.

37. Carlevaro-Fita, J., Rahim, A., Guigo, R., Vardy, L.A., and Johnson, R. (2016). Cytoplasmic long noncoding RNAs are frequently bound to and degraded at ribosomes in human cells. RNA 22, 867–882. 10.1261/rna.053561.115.

38. Ji, P., Diederichs, S., Wang, W., Böing, S., Metzger, R., Schneider, P.M., Tidow, N., Brandt, B., Buerger, H., and Bulk, E. (2003). MALAT-1, a novel noncoding RNA, and thymosin β4 predict metastasis and survival in early-stage non-small cell lung cancer. Oncogene 22, 8031–8041.

39. Weksberg, R., Nishikawa, J., Caluseriu, O., Fei, Y.-L., Shuman, C., Wei, C., Steele, L., Cameron, J., Smith, A., and Ambus, I. (2001). Tumor development in the Beckwith– Wiedemann syndrome is associated with a variety of constitutional molecular 11p15 alterations including imprinting defects of KCNQ1OT1. Human molecular genetics 10, 2989–3000.

40. Wu, Y. (2023). SNHG11: A New Budding Star in Tumors and Inflammatory Diseases. Mini Reviews in Medicinal Chemistry.

41. Zhu, W., and Oxford, G.S. (2011). Differential gene expression of neonatal and adult DRG neurons correlates with the differential sensitization of TRPV1 responses to nerve growth factor. Neuroscience letters 500, 192–196.

42. Bayat, A., Liu, Z., Luo, S., Fenger, C.D., Højte, A.F., Isidor, B., Cogne, B., Larson, A., Zanus, C., and Faletra, F. (2023). A new neurodevelopmental disorder linked to heterozygous variants in UNC79. Genetics in Medicine 25, 100894.

43. Uchida, M., Enomoto, A., Fukuda, T., Kurokawa, K., Maeda, K., Kodama, Y., Asai, N., Hasegawa, T., Shimono, Y., and Jijiwa, M. (2006). Dok-4 regulates GDNF-dependent neurite outgrowth through downstream activation of Rap1 and mitogen-activated protein kinase. Journal of cell science 119, 3067–3077.

44. Müller, M.B., Kasturi, P., Jayaraj, G.G., and Hartl, F.U. (2023). Mechanisms of readthrough mitigation reveal principles of GCN1-mediated translational quality control. Cell.

45. Raftogianis, R.B., Wood, T.C., Otterness, D.M., Van Loon, J.A., and Weinshilboum, R.M. (1997). Phenol sulfotransferase pharmacogenetics in humans: association of commonSULT1A1alleles with TS PST phenotype. Biochemical and biophysical research communications 239, 298–304.

46. Sidharthan, N.P., Minchin, R.F., and Butcher, N.J. (2013). Cytosolic sulfotransferase 1A3 is induced by dopamine and protects neuronal cells from dopamine toxicity: role of D1 receptor-N-methyl-D-aspartate receptor coupling. Journal of Biological Chemistry 288, 34364–34374.

47. Zhou, X., Wang, N., Liu, W., Chen, R., Yang, G., and Yu, H. (2023). Identification of the potential association between SARS-CoV-2 infection and acute kidney injury based on the shared gene signatures and regulatory network. BMC Infectious Diseases 23, 655.

48. Chen, S.-P., Zhou, Y.-Q., Liu, D.-Q., Zhang, W., Manyande, A., Guan, X.-H., Tian, Y.-k., Ye, D.-W., and Mohamed Omar, D. (2017). PI3K/Akt pathway: a potential therapeutic target for chronic pain. Current pharmaceutical design 23, 1860–1868.

50. Bangash, M., Alles, S.R., Santana-Varela, S., Millet, Q., Sikandar, S., de Clauser, L., Ter Heegde, F., Habib, A.M., Pereira, V., and Sexton, J.E. (2018). Distinct transcriptional responses of mouse sensory neurons in models of human chronic pain conditions. Wellcome open research 3.

51. Das, V., Kc, R., Li, X., Varma, D., Qiu, S., Kroin, J.S., Forsyth, C.B., Keshavarzian, A., van Wijnen, A.J., and Park, T.J. (2018). Pharmacological targeting of the mammalian clock reveals a novel analgesic for osteoarthritis-induced pain. Gene 655, 1–12.

51. Dussor, G., Zylka, M.J., Anderson, D.J., and McCleskey, E.W. (2008). Cutaneous sensory neurons expressing the Mrgprd receptor sense extracellular ATP and are putative nociceptors. Journal of neurophysiology 99, 1581–1589.

52. Osada, S.-I., Mizuno, K., Saido, T.C., Suzuki, K., Kuroki, T., and Ohno, S. (1992). A new member of the protein kinase C family, nPKC theta, predominantly expressed in skeletal muscle. Molecular and cellular biology.

53. Kusuda, J., Hirai, M., Tanuma, R., and Hashimoto, K. (2000). Cloning, expression analysis and chromosome mapping of human casein kinase 1 γ1 (CSNK1G1): Identification of two types of cDNA encoding the kinase protein associated with heterologous carboxy-terminal sequences. Cytogenetics and cell genetics 90, 298–302.

54. Ziembik, M.A., Bender, T.P., Larner, J.M., and Brautigan, D.L. (2017). Functions of protein phosphatase-6 in NF-κB signaling and in lymphocytes. Biochemical Society Transactions 45, 693–701.

55. Jeronimo, C., Forget, D., Bouchard, A., Li, Q., Chua, G., Poitras, C., Thérien, C., Bergeron, D., Bourassa, S., and Greenblatt, J. (2007). Systematic analysis of the protein interaction network for the human transcription machinery reveals the identity of the 7SK capping enzyme. Molecular cell 27, 262–274.

56. Bergfort, A., Hilal, T., Kuropka, B., Ilik, İ.A., Weber, G., Aktaş, T., Freund, C., and Wahl, M.C. (2022). The intrinsically disordered TSSC4 protein acts as a helicase inhibitor, placeholder and multi-interaction coordinator during snRNP assembly and recycling. Nucleic Acids Research 50, 2938–2958.

57. Cottle, W.T., Wallert, C.H., Anderson, K.K., Tran, M.F., Bakker, C.L., Wallert, M.A., and Provost, J.J. (2020). Calcineurin homologous protein isoform 2 supports tumor survival via the sodium hydrogen exchanger isoform 1 in non-small cell lung cancer. Tumor Biology 42, 1010428320937863.

58. Holtz, A.M., Griffiths, S.C., Davis, S.J., Bishop, B., Siebold, C., and Allen, B.L. (2015). Secreted HHIP1 interacts with heparan sulfate and regulates Hedgehog ligand localization and function. Journal of Cell Biology 209, 739–758.

59. Xing, R., Zhou, H., Jian, Y., Li, L., Wang, M., Liu, N., Yin, Q., Liang, Z., Guo, W., and Yang, C. (2021). The Rab7 effector WDR91 promotes autophagy-lysosome degradation in neurons by regulating lysosome fusion. Journal of Cell Biology 220, e202007061.

60. Grootjans, J.J., Zimmermann, P., Reekmans, G., Smets, A., Degeest, G., Dürr, J., and David, G. (1997). Syntenin, a PDZ protein that binds syndecan cytoplasmic domains. Proceedings of the National Academy of Sciences 94, 13683–13688.

61. Berman, D.M., Kozasa, T., and Gilman, A.G. (1996). The GTPase-activating protein RGS4 stabilizes the transition state for nucleotide hydrolysis. Journal of Biological Chemistry 271, 27209–27212.

62. Canela, L., Luján, R., Lluís, C., Burgueño, J., Mallol, J., Canela, E.I., Franco, R., and Ciruela, F. (2007). The neuronal Ca2+-binding protein 2 (NECAB2) interacts with the adenosine A2A receptor and modulates the cell surface expression and function of the receptor. Molecular and Cellular Neuroscience 36, 1–12.

63. Chen, Z., Long, H., Guo, J., Wang, Y., He, K., Tao, C., Li, X., Jiang, K., Guo, S., and Pi, Y. (2022). Autism-Risk Gene necab2 Regulates Psychomotor and Social Behavior as a Neuronal Modulator of mGluR1 Signaling. Frontiers in Molecular Neuroscience 15, 901682.

64. Nalamalapu, R.R., Yue, M., Stone, A.R., Murphy, S., and Saha, M.S. (2021). The tweety gene family: from embryo to disease. Frontiers in Molecular Neuroscience 14, 672511.

65. Boulpicante, M., Darrigrand, R., Pierson, A., Salgues, V., Rouillon, M., Gaudineau, B., Khaled, M., Cattaneo, A., Bachi, A., and Cascio, P. (2020). Tumors escape immunosurveillance by overexpressing the proteasome activator PSME3. Oncoimmunology 9, 1761205.

66. Wang, Z., Zhang, M., Shan, R., Wang, Y.J., Chen, J., Huang, J., Sun, L.Q., and Zhou, W.B. (2019). MTMR3 is upregulated in patients with breast cancer and regulates proliferation, cell cycle progression and autophagy in breast cancer cells. Oncology Reports 42, 1915–1923.

67. Hao, S.-W., Li, T.-R., Han, C., Han, Y., and Cai, Y.-N. (2023). Associations Between Levels of Peripheral NCAPH2 Promoter Methylation and Different Stages of Alzheimer’s Disease: A Cross-Sectional Study. Journal of Alzheimer’s Disease, 1-11.

68. Sander, E., and Collard, J. (1999). Rho-like GTPases: their role in epithelial cell–cell adhesion and invasion. European Journal of Cancer 35, 1905–1911.

69. McEwan, D.G., and Ryan, K.M. (2022). ATG2 and VPS13 proteins: molecular highways transporting lipids to drive membrane expansion and organelle communication. The FEBS Journal 289, 7113–7127.

70. Lauri, S.E., Braithwaite, S.P., Oise Coussen, F., Mulle, C., Dev, K.K., Coutinho, V., Meyer, G., Isaac, J.T., Collingridge, G.L., and Henley, J.M. (2003). Rapid and Differential Regulation of AMPA and Kainate Receptors at Hippocampal Mossy Fibre Synapses by PICK1 and GRIP. Neuron 38, 673.

71. Pengue, G., Cannada-Bartoli, P., and Lania, L. (1993). The ZNF35 human zinc finger gene encodes a sequence-specific DNA-binding protein. FEBS letters 321, 233–236.

72. Merienne, K., Jacquot, S., Pannetier, S., Zeniou, M., Bankier, A., Gecz, J., Mandel, J.-L., Mulley, J., Sassone-Corsi, P., and Hanauer, A. (1999). A missense mutation in RPS6KA3 (RSK2) responsible for non-specific mental retardation. Nature genetics 22, 13–14.

73. Kumar, R., Palmer, E., Gardner, A.E., Carroll, R., Banka, S., Abdelhadi, O., Donnai, D., Elgersma, Y., Curry, C.J., and Gardham, A. (2020). Expanding clinical presentations due to variations in THOC2 mRNA nuclear export factor. Frontiers in Molecular Neuroscience 13, 12.

75. Kloft, N., Neukirch, C., von Hoven, G., Bobkiewicz, W., Weis, S., Boller, K., and Husmann, M. (2012). A subunit of eukaryotic translation initiation factor 2α-phosphatase (CreP/PPP1R15B) regulates membrane traffic. Journal of Biological Chemistry 287, 35299–35317.

75. Escamez, T., Bahamonde, O., Tabares-Seisdedos, R., Vieta, E., Martinez, S., and Echevarria, D. (2012). Developmental dynamics of PAFAH1B subunits during mouse brain development. Journal of Comparative Neurology 520, 3877–3894.

76. Gasparski, A.N., Mason, D.E., Moissoglu, K., and Mili, S. (2022). Regulation and outcomes of localized RNA translation. Wiley Interdisciplinary Reviews: RNA 13, e1721.

77. Jackson, R.J., Hellen, C.U., and Pestova, T.V. (2010). The mechanism of eukaryotic translation initiation and principles of its regulation. Nat Rev Mol Cell Biol 11, 113–127. 10.1038/nrm2838.

78. Kurihara, Y., Matsui, A., Hanada, K., Kawashima, M., Ishida, J., Morosawa, T., Tanaka, M., Kaminuma, E., Mochizuki, Y., Matsushima, A., et al. (2009). Genome-wide suppression of aberrant mRNA-like noncoding RNAs by NMD in Arabidopsis. Proc Natl Acad Sci U S A 106, 2453–2458. 10.1073/pnas.0808902106.

79. Minati, L., Firrito, C., Del Piano, A., Peretti, A., Sidoli, S., Peroni, D., Belli, R., Gandolfi, F., Romanel, A., Bernabo, P., et al. (2021). One-shot analysis of translated mammalian lncRNAs with AHARIBO. Elife 10. 10.7554/eLife.59303.

80. Anderson, D.M., Anderson, K.M., Chang, C.L., Makarewich, C.A., Nelson, B.R., McAnally, J.R., Kasaragod, P., Shelton, J.M., Liou, J., Bassel-Duby, R., and Olson, E.N. (2015). A micropeptide encoded by a putative long noncoding RNA regulates muscle performance. Cell 160, 595–606. 10.1016/j.cell.2015.01.009.

81. Ruiz-Orera, J., Messeguer, X., Subirana, J.A., and Alba, M.M. (2014). Long non-coding RNAs as a source of new peptides. Elife 3, e03523. 10.7554/eLife.03523.

82. Carrieri, C., Cimatti, L., Biagioli, M., Beugnet, A., Zucchelli, S., Fedele, S., Pesce, E., Ferrer, I., Collavin, L., Santoro, C., et al. (2012). Long non-coding antisense RNA controls Uchl1 translation through an embedded SINEB2 repeat. Nature 491, 454–457. 10.1038/nature11508.

83. Yoon, J.H., Abdelmohsen, K., Srikantan, S., Yang, X., Martindale, J.L., De, S., Huarte, M., Zhan, M., Becker, K.G., and Gorospe, M. (2012). LincRNA-p21 suppresses target mRNA translation. Mol Cell 47, 648–655. 10.1016/j.molcel.2012.06.027.

84. Pecoraro, V., Rosina, A., and Polacek, N. (2022). Ribosome-Associated ncRNAs (rancRNAs) Adjust Translation and Shape Proteomes. Noncoding RNA 8. 10.3390/ncrna8020022.

85. Pircher, A., Gebetsberger, J., and Polacek, N. (2014). Ribosome-associated ncRNAs: an emerging class of translation regulators. RNA Biol 11, 1335–1339. 10.1080/15476286.2014.996459.

86. Sapio, M.R., Iadarola, M.J., Loydpierson, A.J., Kim, J.J., Thierry-Mieg, D., Thierry-Mieg, J., Maric, D., and Mannes, A.J. (2020). Dynorphin and enkephalin opioid peptides and transcripts in spinal cord and dorsal root ganglion during peripheral inflammatory hyperalgesia and allodynia. The Journal of pain 21, 988–1004.

88. Prochiantz, A., and Di Nardo, A.A. (2022). Shuttling Homeoproteins and Their Biological Significance. Cell Penetrating Peptides: Methods and Protocols, 33-44.

88. Catela, C., Chen, Y., Weng, Y., Wen, K., and Kratsios, P. (2022). Control of spinal motor neuron terminal differentiation through sustained Hoxc8 gene activity. Elife 11, e70766.

89. Heiman, M., Kulicke, R., Fenster, R.J., Greengard, P., and Heintz, N. (2014). Cell type-specific mRNA purification by translating ribosome affinity purification (TRAP). Nat Protoc 9, 1282–1291. 10.1038/nprot.2014.085.

90. Heiman, M., Kulicke, R., Fenster, R.J., Greengard, P., and Heintz, N. (2014). Cell type– specific mRNA purification by translating ribosome affinity purification (TRAP). Nature protocols 9, 1282–1291.

91. Kanellopoulos, A.H., Koenig, J., Huang, H., Pyrski, M., Millet, Q., Lolignier, S., Morohashi, T., Gossage, S.J., Jay, M., Linley, J.E., et al. (2018). Mapping protein interactions of sodium channel NaV1.7 using epitope-tagged gene-targeted mice. EMBO J 37, 427–445. 10.15252/embj.201796692.

92. Ray, P.R., Shiers, S., Caruso, J.P., Tavares-Ferreira, D., Sankaranarayanan, I., Uhelski, M.L., Li, Y., North, R.Y., Tatsui, C., and Dussor, G. (2023). RNA profiling of human dorsal root ganglia reveals sex differences in mechanisms promoting neuropathic pain. Brain 146, 749–766.

94. Sierra, C., De Toma, I., and Dierssen, M. (2021). Single nucleus RNA-seq in the hippocampus of a Down syndrome mouse model reveals new key players in memory. bioRxiv, 2021.2011. 2018.469102.

94. Minett, M.S., Quick, K., and Wood, J.N. (2011). Behavioral measures of pain thresholds. Current protocols in mouse biology 1, 383–412.

95. Haroun, R., Gossage, S.J., Luiz, A.P., Arcangeletti, M., Sikandar, S., Zhao, J., Cox, J.J., and Wood, J.N. (2023). Chemogenetic Silencing of NaV1. 8-Positive Sensory Neurons Reverses Chronic Neuropathic and Bone Cancer Pain in FLEx PSAM4-GlyR Mice. eneuro 10.

96. Randall, L.O. (1957). A method for measurement of analgesic activity of inflamed tissue. Arch Int Pharmacodyn Ther 111, 409–411.

97. Chaplan, S.R., Bach, F.W., Pogrel, J., Chung, J., and Yaksh, T. (1994). Quantitative assessment of tactile allodynia in the rat paw. Journal of neuroscience methods 53, 55–63.

98. Hargreaves, K., Dubner, R., Brown, F., Flores, C., and Joris, J. (1988). A new and sensitive method for measuring thermal nociception in cutaneous hyperalgesia. Pain 32, 77–88.

