## Supplemental Files for "Analgesic targets identified in mouse sensory neuron somata and terminal pain translatomes"

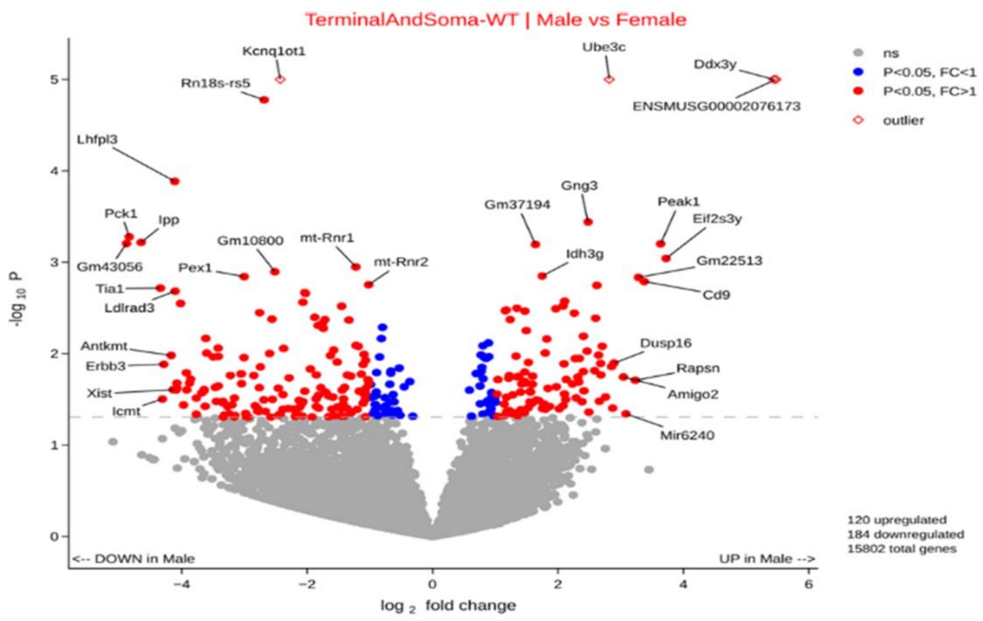


**Figure S1 (related to Figure 2):** Volcano plot showing differentially expressed translated genes in both somata and terminals between male (n=6) and female (n=6) mouse samples. Genes on the left-hand side of the plot are upregulated in females and genes upregulated in males are shown on the right-hand side. Non-significant transcripts are grey, significant genes ($p<0.05$) with $\log_2$ fold change < 1 are marked in blue and significant genes with $\log_2$ fold change > 1 are represented with a red colour. X-axis and Y-axis show, $\log_2$ fold change and $-\log_{10}$ p-value, respectively. Complete data comprising read numbers, fold increase ($\log_2$) and p-values are presented in tables S1 and S2.

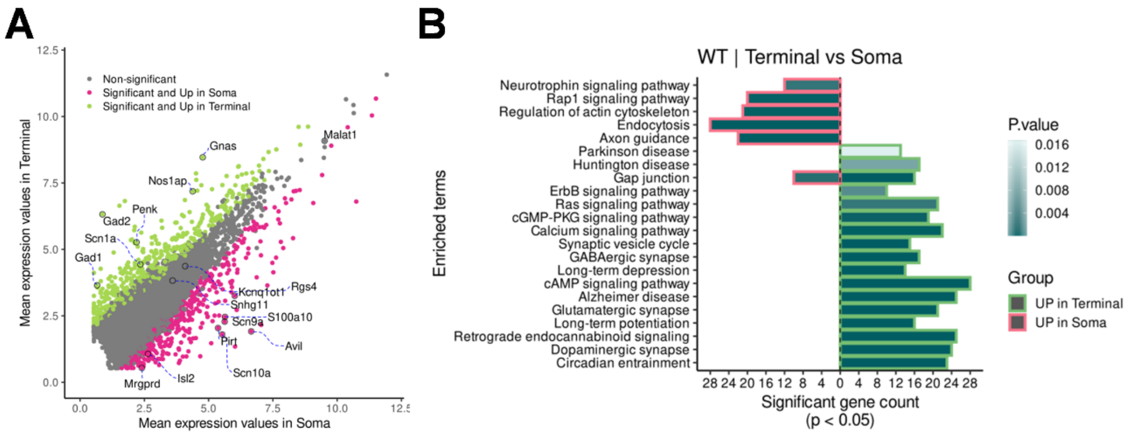


**Figure S2 (related to Figure 3): Soma vs Terminal further comparisons.**

**A** Scatter Plot showing the comparison of the mean expression values of all genes between the soma and terminal of wildtype mice. Non-significant transcripts are grey, significant genes ($p<0.05$) upregulated in the soma are pink and significant genes ($p<0.05$) upregulated in the terminals are represented with a green colour. X-axis and Y-axis show mean expression values in soma and terminals, respectively.

**B** Horizontal bar plot showing the different signalling pathways of the genes enriched in the terminal (green) or soma (red). X-axis shows the number of genes significantly enriched in the two conditions, while Y-axis shows the pathways they refer to.

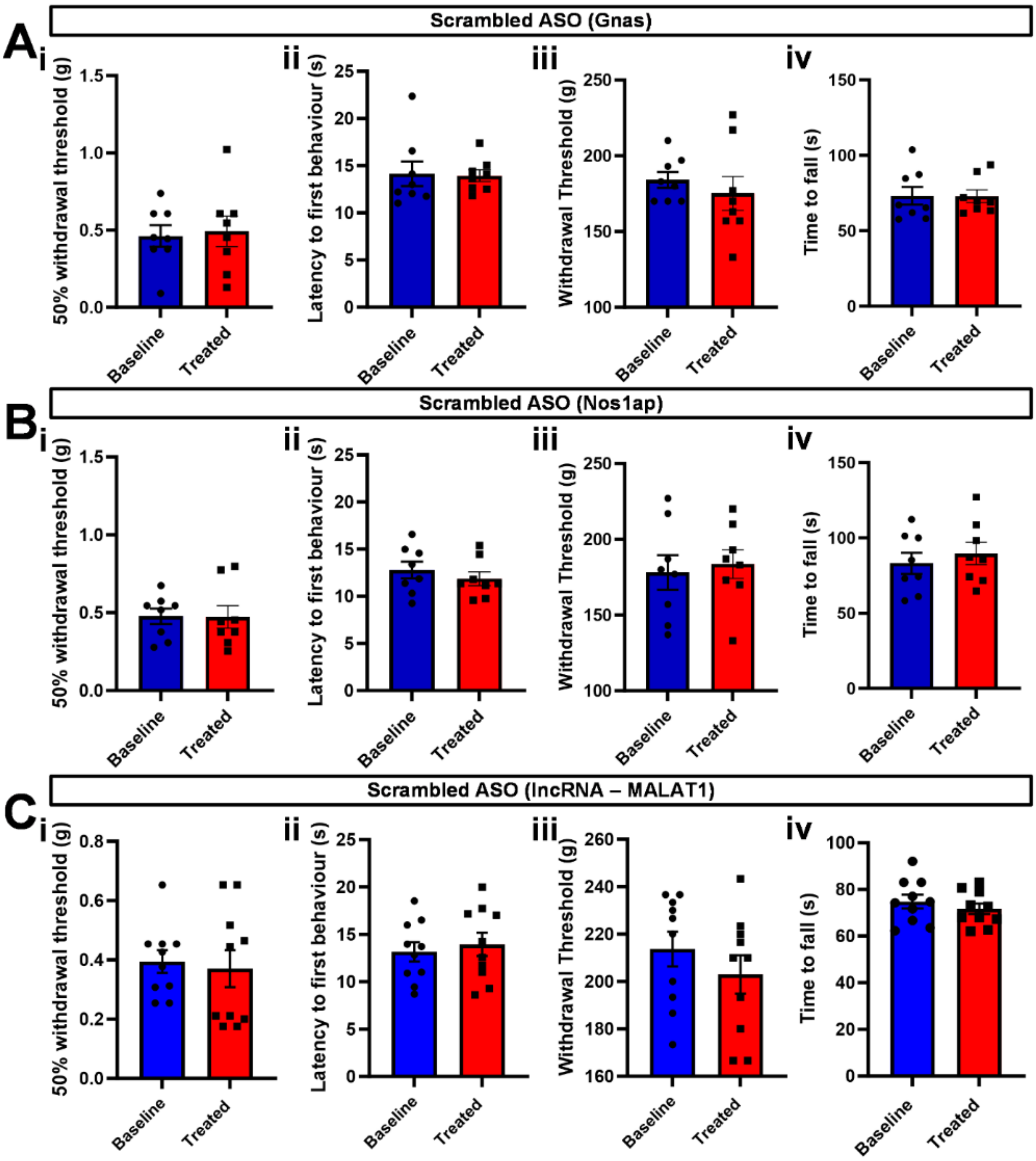

Scrambled ASO (Gnas)
A
i
50% withdrawal threshold (g)
ii
Latency to first behaviour (s)
iii
Withdrawal Threshold (g)
iv
Time to fall (s)
Baseline
Treated
Scrambled ASO (Nos1ap)
B
Scrambled ASO (lncRNA – MALAT1)
C

**Figure S3 (related to Figure 3): Control antisense oligonucleotide pain behavioural tests.**

**A** Acute behavioural tests battery showing no effect of the control (scrambled) ASOs designed based on the Gnas ASOs on mechanical and thermal thresholds as well as motor coordination (**i**: Von Frey, **ii**:Hargreaves, **iii**: Randall-Selitto, **iv**: Rotarod).
**B** Acute behavioural tests battery showing no effect of the control (scrambled) ASOs designed based on the Nos1ap ASOs on mechanical and thermal thresholds as well as motor coordination (**i**: Von Frey, **ii**:Hargreaves, **iii**: Randall-Selitto, **iv**: Rotarod).
**C** Acute behavioural tests battery showing no effect of the control (scrambled) ASOs designed based on the ASOs anti lncRNA (MALAT1) on mechanical and thermal thresholds as well as motor coordination (**i**: Von Frey, **ii**:Hargreaves, **iii**: Randall-Selitto, **iv**: Rotarod).
Data are represented as mean ± SEM. Mean latencies, withdrawal thresholds, and times to fall were compared using paired t-test.

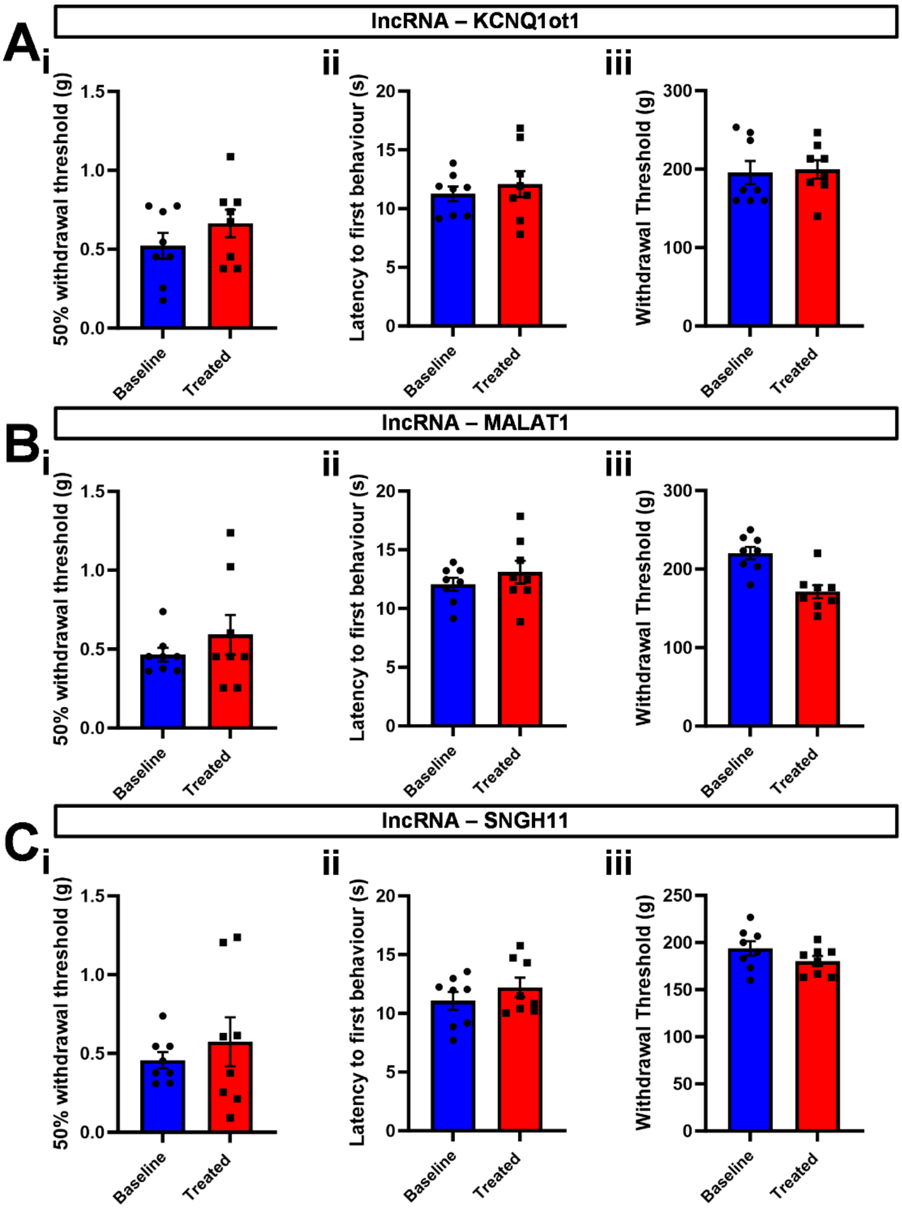
A
lncRNA – KCNQ1ot1
i
ii
iii
50% withdrawal threshold (g)
Latency to first behaviour (s)
Withdrawal Threshold (g)
Baseline
Treated
B
lncRNA – MALAT1
C
lncRNA – SNGH11

**Figure S4 (related to Figure 3): Effect of single long non-coding RNA ASOs on acute pain.**

**A** Mechanical and thermal acute pain thresholds assessed using Von Frey **(i)**, Hargreaves **(ii)**, and Randall-Selitto tests **(iii)** to mice before (blue) and after (red) treatment with ASO targeted to KNCQ11ot1.

**B** Mechanical and thermal acute pain thresholds assessed using Von Frey **(i)**, Hargreaves **(ii)**, and Randall-Selitto tests **(iii)** to mice before (blue) and after (red) treatment with ASO targeted to MALAT1.

**C** Mechanical and thermal acute pain thresholds assessed using Von Frey **(i)**, Hargreaves **(ii)**, and Randall-Selitto tests **(iii)** to mice before (blue) and after (red) treatment with ASO targeted to SNHG11.

Data are represented as mean ± SEM. Mean latencies and withdrawal thresholds were compared using paired t-test.

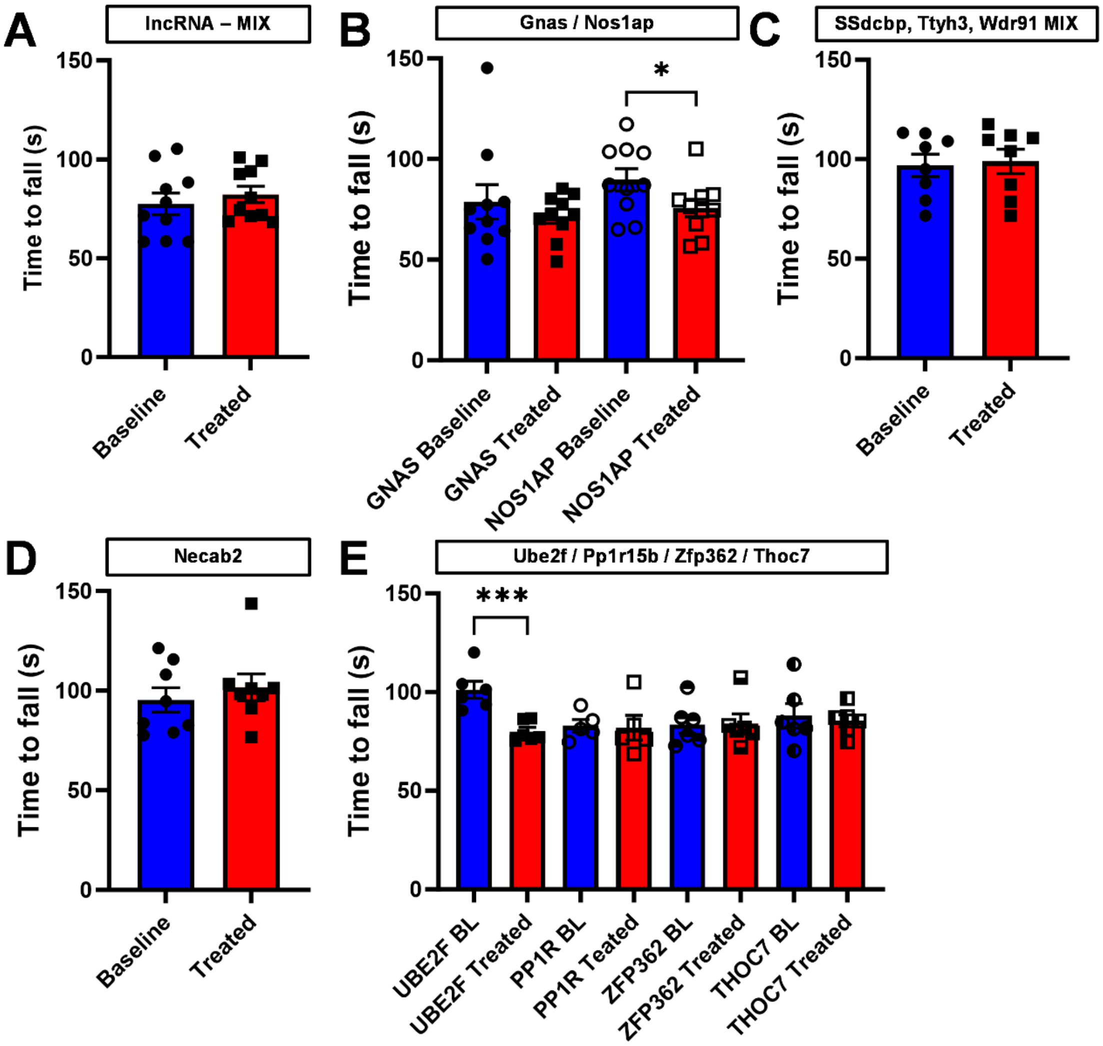

A
lncRNA – MIX
Time to fall (s)
150
100
50
0
Baseline
Treated
B
Gnas / Nos1ap
*
GNAS Baseline
GNAS Treated
NOS1AP Baseline
NOS1AP Treated
C
SSdcbp, Ttyh3, Wdr91 MIX
D
Necab2
E
Ube2f / Pp1r15b / Zfp362 / Thoc7
***
UBE2F BL
UBE2F Treated
PP1R BL
PP1R Teated
ZFP362 BL
ZFP362 Treated
THOC7 BL
THOC7 Treated

**Figure S5 (related to Figures 3, 7, 8): Motor coordination tests in mice treated with ASOs.**

**A** Motor coordination assessment using Rotarod for mice treated with ASO targeted towards a mix of long non-coding RNAs (KCNQ11OT1, MALAT1, SNHG11).

**B** Motor coordination assessment using Rotarod for mice treated with ASO targeted towards GNAS and Nos1AP.

**C** Motor coordination assessment using Rotarod for mice treated with ASO targeted towards a mix of genes enriched in the soma of mice treated with NGF (Sdcbp, Ttyh3, Wdr91).

**D** Motor coordination assessment using Rotarod for mice treated with ASO targeted towards NECAB2.

**E** Motor coordination assessment using Rotarod for mice treated with ASO targeted towards a battery of four genes enriched in the terminal of mice treated with NGF (UBE2F, Ppp1r15b, Zfp362, Thoc7).

Data are represented as mean ± SEM. Times to fall were compared using paired t-test. * = $p<0.05$. *** = $p<0.001$.

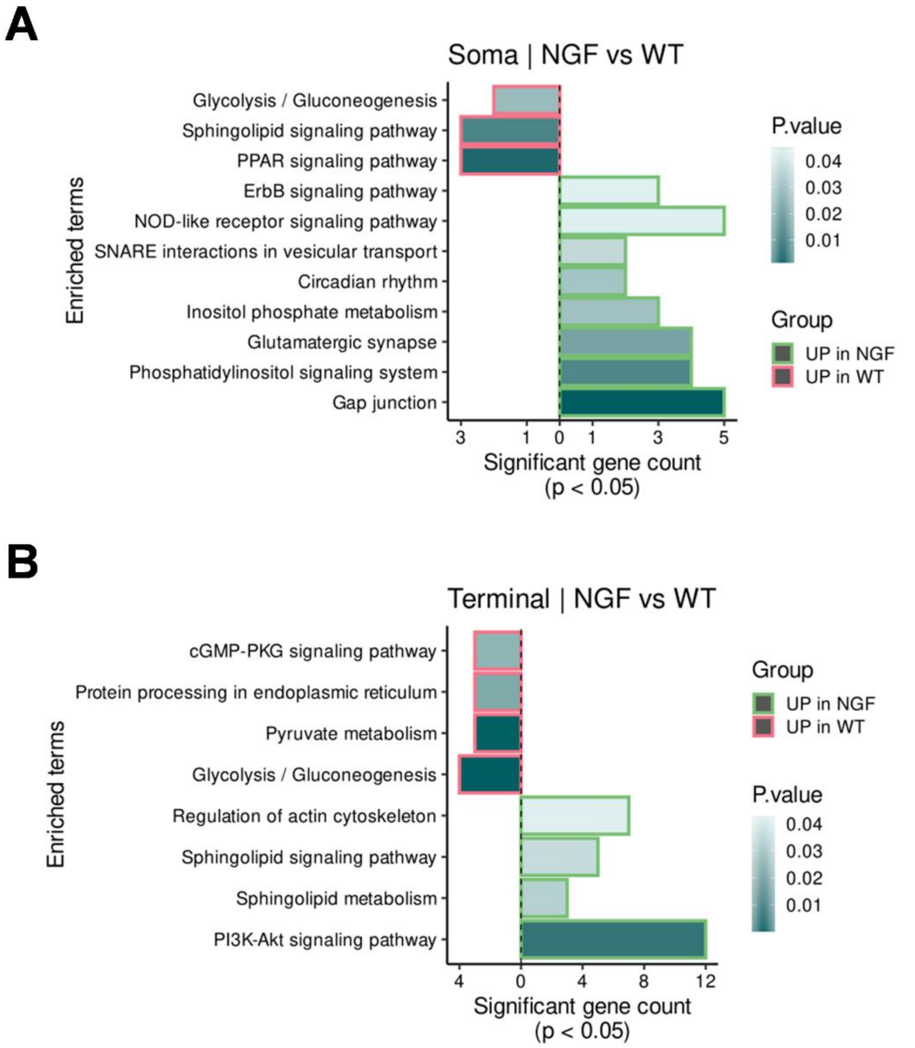


**Figure S6 (related to Figure 4): Enriched gene pathways of NGF-treated versus wildtype mice.**

**A** Horizontal bar plots showing the different signalling pathways of the genes enriched in the somata of NGF-treated (green) or wildtype mice (red).

**B** Horizontal bar plots showing the different signalling pathways of the genes enriched in the terminal of NGF-treated (green) or wildtype mice(red). X-axis shows the number of genes significantly enriched in the two conditions, while Y-axis shows the pathways they refer to.

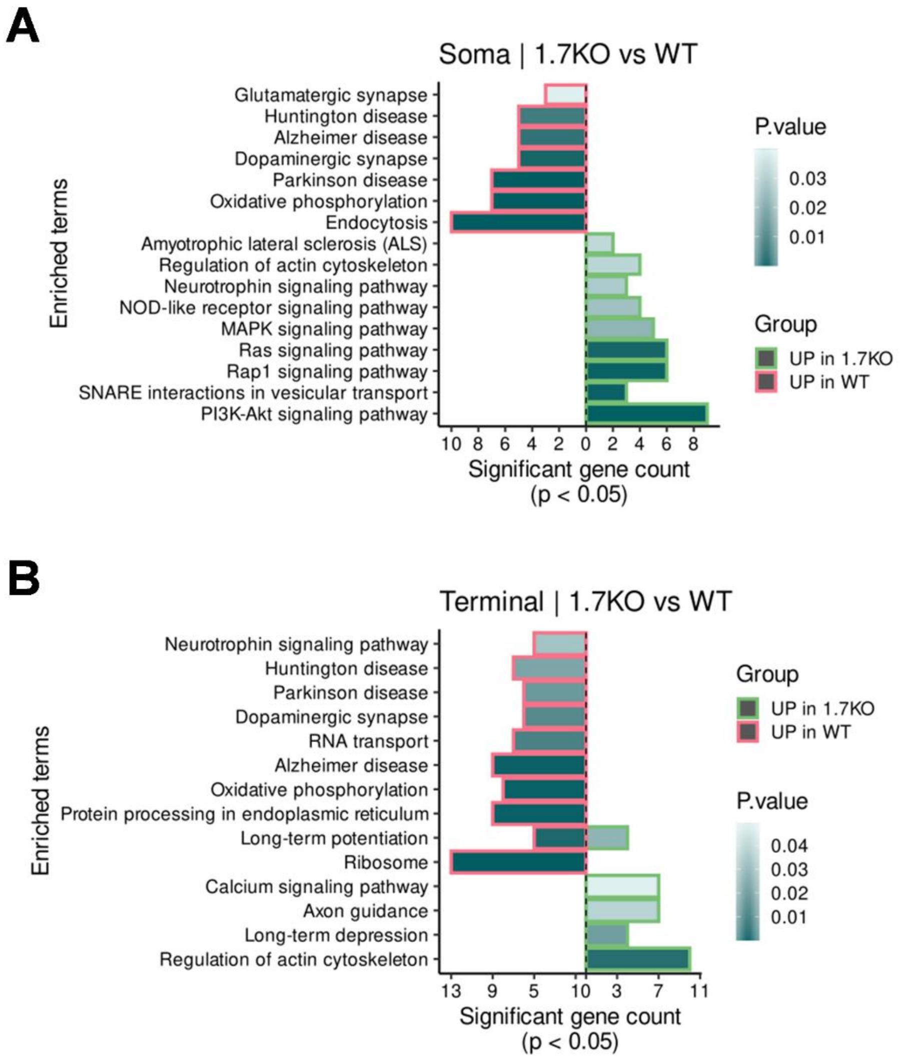


**Figure S7 (related to Figure 5): Enriched gene pathways of NaV1.7-KO versus wildtype mice.**

**A** Horizontal bar plots showing the different signalling pathways of the genes enriched in the somata of NaV1.7-KO (green) or wildtype mice (red).

**B** Horizontal bar plots showing the different signalling pathways of the genes enriched in the terminal of NaV1.7-KO (green) or wildtype mice (red). X-axis shows the number of genes significantly enriched in the two conditions, while Y-axis shows the pathways they refer to.

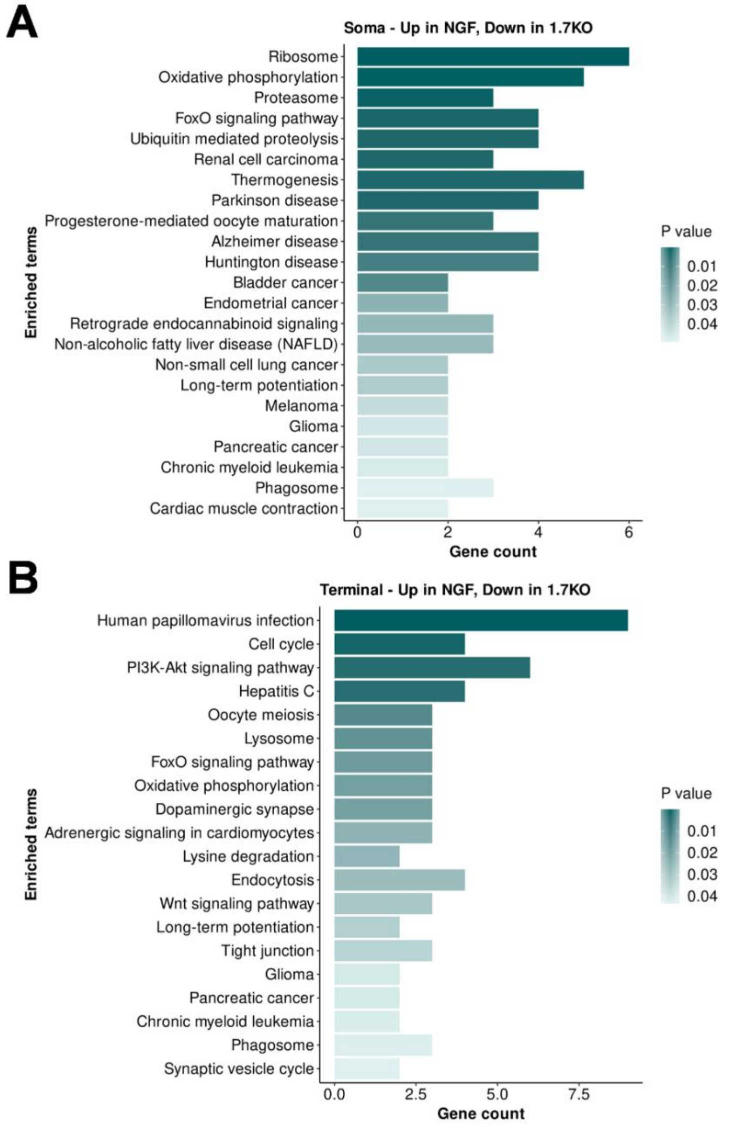


**Figure S8 (related to Figure 6): Enriched gene pathways of NGF-treated versus NaV1.7-KO mice.**

**A)** Horizontal bar plots showing the different signalling pathways of the genes up-regulated in the somata of NGF-treated mice and down-regulated in NaV1.7-KO mice.

**B)** Horizontal bar plots showing the different signalling pathways of the genes up-regulated in the terminals of NGF-treated mice and down-regulated in NaV1.7-KO mice. X-axis shows the number of genes significantly enriched in the two conditions, while Y-axis shows the pathways they refer to.

### Supplemental Files

**Table S1: RNA-Seq data from somata of Males vs Female mice.**

Read counts, fold change, and p-values are found in this table.

**Table S2: RNA-Seq data from terminals of Males vs Female mice.**

Read counts, fold change, and p-values are found in this table.

**Table S3: RNA-Seq data from Somata vs Terminal samples in mice.**

Read counts, fold change, and p-values are found in this table.

**Table S4: RNA-Seq data from somata of NGF-treated vs WT mice.**

Read counts, fold change, and p-values are found in this table.

**Table S5: RNA-Seq data from terminals of NGF-treated vs WT mice.**

Read counts, fold change, and p-values are found in this table.

**Table S6: RNA-Seq data from somata of NaV1.7 null vs WT mice.**

Read counts, fold change, and p-values are found in this table.

**Table S7: RNA-Seq data from terminals of NaV1.7 null vs WT mice.**

Read counts, fold change, and p-values are found in this table.

**Table S8: RNA-Seq data from somata of NaV1.7 null vs NGF-treated mice.**

Read counts, fold change, and p-values are found in this table.

**Table S9: RNA-Seq data from somata of NaV1.7 null vs NGF-treated mice.**

Read counts, fold change, and p-values are found in this table.
